# Lack of dynamic balance between vesicle transport and microtubule growth in neurite outgrowth enables formation of dystrophic bulbs

**DOI:** 10.1101/153569

**Authors:** Arjun Singh Yadaw, Mustafa M. Siddiq, Vera Rabinovich, Rosa Tolentino, Ravi Iyengar, Jens Hansen

## Abstract

Whole cell responses involve multiple subcellular processes (SCPs). To understand how balance between SCPs controls the dynamics of whole cell responses we studied neurite outgrowth in rat primary cortical neurons in culture. We used a combination of dynamical models and experiments to understand the conditions that permitted growth at a specified velocity and when aberrant growth could lead to the formation of dystrophic bulbs. We hypothesized that dystrophic bulb formation is due to quantitative imbalances between SCPs. Simulations predict redundancies between lower level sibling SCPs within each type of high level SCP. In contrast, higher level SCPs, such as vesicle transport and exocytosis or microtubule growth characteristic of each type need to be strictly coordinated with each other and imbalances result in stalling of neurite outgrowth. From these simulations, we predicted the effect of changing the activities of SCPs involved in vesicle exocytosis or microtubule growth could lead to formation of dystrophic bulbs. siRNA ablation experiments verified these predictions. We conclude that whole cell dynamics requires balance between the higher-level SCPs involved and imbalances can terminate whole cell responses such as neurite outgrowth.

## Introduction

Neurite outgrowth is an early event that changes the state of neurons (Arimura and Kaibuchi, 2007) and allows neurons to develop axons and dendritic trees that connect to other neurons and become electrically active. When there is nerve injury and the axons are severed, the process of regeneration which is similar to neurite outgrowth often fails, resulting in the formation of bulbs at the ends of regenerating axons. These bulbs are called dystrophic bulbs (Hill, 2017). Although some of the cellular pathways involved in axonal regeneration, failure and formation of the dystrophic bulbs are known (Blanquie and Bradke, 2018; Filious and Schwab, 2018) the mechanisms by which these lead to failure are not well understood. In this study we have used an integrated computational and experimental approach to understand the origins of dystrophic bulbs and more generally the conditions under which whole cell responses such as neurite outgrowth can be maintained. The central question we asked in this study is whether quantitative imbalances between subcellular processes can lead to the formation of dystrophic bulbs.

Whole cell responses that involve, both morphological and physiological changes, are complex because they can engage many subcellular processes (SCPs) that are together responsible for whole cell functions. SCPs include biochemical pathways and cell biological processes involved the functions of different organelles. Coordinated changes in the activities of SCPs often lead to a change in cell state, such as moving to a more differentiated phenotype. Detailed understanding of how different SCPs collaborate to produce an integrated whole cell response requires experiments, along with mathematical models that capture the dynamics of individual SCPs as well as the interrelationship between the dynamics of multiple SCPs. Based on the biological detail that is represented by the SCPs they can be classified as Level-1, Level-2 and Level-3 SCPs. Higher level SCPs (i.e. SCPs with smaller numbers) represent a cell biological function that combines the functions of their lower level SCP children (Hansen et al., 2017). To understand whole cell responses, it is necessary to delineate the quantitative coordination between the dynamics of the SCPs. In the case of neurite outgrowth, three Level-1 SCPs, *Production of Membrane Components, Vesicle Transport & Exocytosis* and *Microtubule Growth* need to be coordinated. Membrane components such as membrane lipids and proteins are synthesized in the cell body and transported to the growth cone via vesicle transport. Vesicles that fuse with the growth cone plasma membrane deliver their membrane lipids to the neurite tip, causing the neurite shaft to grow. Microtubules form the basis of the neurite scaffold that needs to extent in coordination with the neurite shaft. Tsaneva-Atanasova et al (Tsaneva-Atanasova et al., 2009) developed a model of the interaction between microtubules and membrane vesicles in regulating neurite outgrowth. The framework of this model serves as a useful starting point for more extensive models where we can consider how a group of distinct but related SCPs such as *Vesicle Budding, Vesicle Transport* and *Vesicle Fusion* along with a second group of SCPs that involves *Microtubule Nucleation, Microtubule Growth* and *Conversion of Dynamic to Stable Microtubules* interact to enable the growth of neurites. Such detailed models could serve as basis for understanding how a single network can regulate the relationship between kinetic parameters and changes in the levels or activities of molecular components within the different SCPs. To develop a detailed model of neurite outgrowth based on interactions between SCPs of different levels, we constructed a multi compartment ordinary differential equation model to extend and integrate the microtubule growth model developed by Margolin et al. (Margolin et al., 2012) with the vesicle transport model by Heinrich and Rapoport (Heinrich and Rapoport, 2005). We have also incorporated more recent experimental data to make the model contemporary.

Many mathematical models of neurite outgrowth have been developed (Kiddie et al., 2005; Wissner-Gross et al., 2011) and these models have provided valuable insights into how complex biological processes can be modeled at various levels of description. Often these models abstract the details of the underlying mechanisms and consequently it is difficult to decipher how the balance between the various SCPs is achieved to generate whole cell responses. Some studies have focused on specific facets of the neurite outgrowth process such the role of signaling and regulatory motifs in determining how particular neurites are selected to become axons (Fivaz et al., 2008). Such models are useful in understanding the regulation of complex cellular responses. In developing our dynamical model of neurite outgrowth, we have made a critical change from our previous approaches to dynamical modeling in developing this model. We had typically used a bottom-up approach where individual reactions are assembled into modules that have emergent behaviors (Azeloglu et al., 2014; Bhalla and Iyengar, 1999). In contrast, the approach we use here can be called top-down as we made a starting postulate that the three Level-1 SCPs involved in neurite outgrowth need to be appropriately coordinated to produce a steady velocity of neurite outgrowth. We start with experimental observations of neurite outgrowth velocities and use numerical simulations to determine how their Level-2 children and Level-3 grandchildren SCPs need to be coordinated to allow neurite outgrowth with the observed velocities and under kinetic parameter constellations that match additional experimental data. Our simulations show redundancy between Level-3 sibling SCP activities. Changes in function of one Level-3 sibling SCP can be compensated by alteration of function in another. However, such generalized reciprocal relationships do not exist between Level-2 SCPs. Change in activity in one Level-2 SCP cannot be simply compensated by activity adaptation of one of its sibling Level-2 SCPs but requires adaptations of multiple sibling SCPs. Consequently, loss of coordinated whole cell function is more likely. We used the findings from analyses of the computational models to develop predictions regarding effects of modulating higher SCPs on whole cell function. We tested the predictions experimentally and found them to be valid.

## Results

### Model Description: Integration of subcellular processes involved in vesicle production, transport and exocytosis and microtubule dynamics to model neurite outgrowth

Neurite outgrowth and axonal regeneration share common SCPs such as vesicle production & transport and microtubule growth in an orderly fashion leading to a growing shaft and a growth cone at the tip as shown schematically in Figure 1A left panel. Failure of growth and the subsequent formation of dystrophic bulb (Figure 1A right panel) can be hypothesized as failure of SCPs to function “properly” with respect to each other. To understand how this lack of coordination between SCPs occurs it is necessary to have a detailed description of the SCPs and their relationships as part of setting up the computational model we provide a detailed description of SCPs and their relationships. Among the major SCPs that are involved in neurite outgrowth (NOG) are the Level-1 SCPs *Production of Membrane Components*, *Vesicle Transport and Exocytosis* and *Microtubule Growth* (Figure 1B). Each of these SCPs is composed of multiple Level-2 children SCPs whose activities need to be coordinated to ensure NOG without complication. Since our major focus was on vesicle transport and microtubule growth, we further categorized Level-2 SCP function by introducing their Level-3 children SCP. SCPs with the same parent SCPs are defined to be siblings. This limited scope enabled us to develop detailed, yet manageable computational models. It is important to note that at this stage our model does not consider local actin dynamics at the growth cone, since we particularly focus on SCPs that are involved in extension of the neurite shaft; i.e. membrane lipid and protein synthesis and delivery and microtubule scaffold growth.

**Figure 1:**
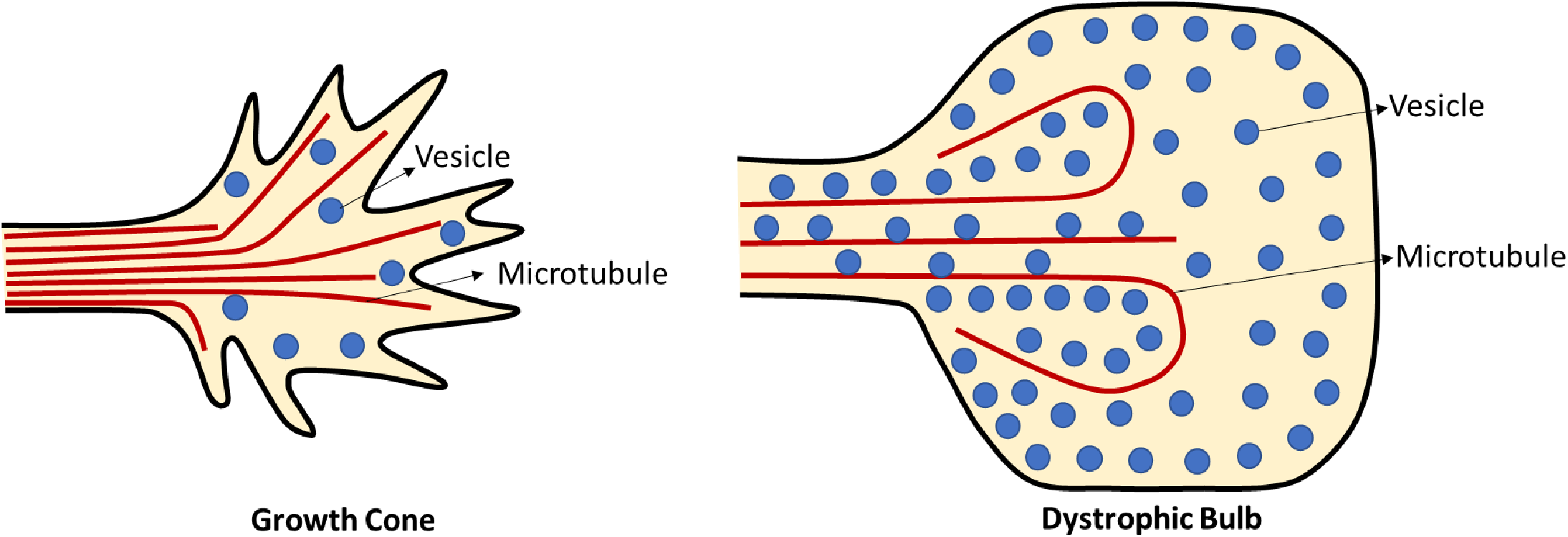
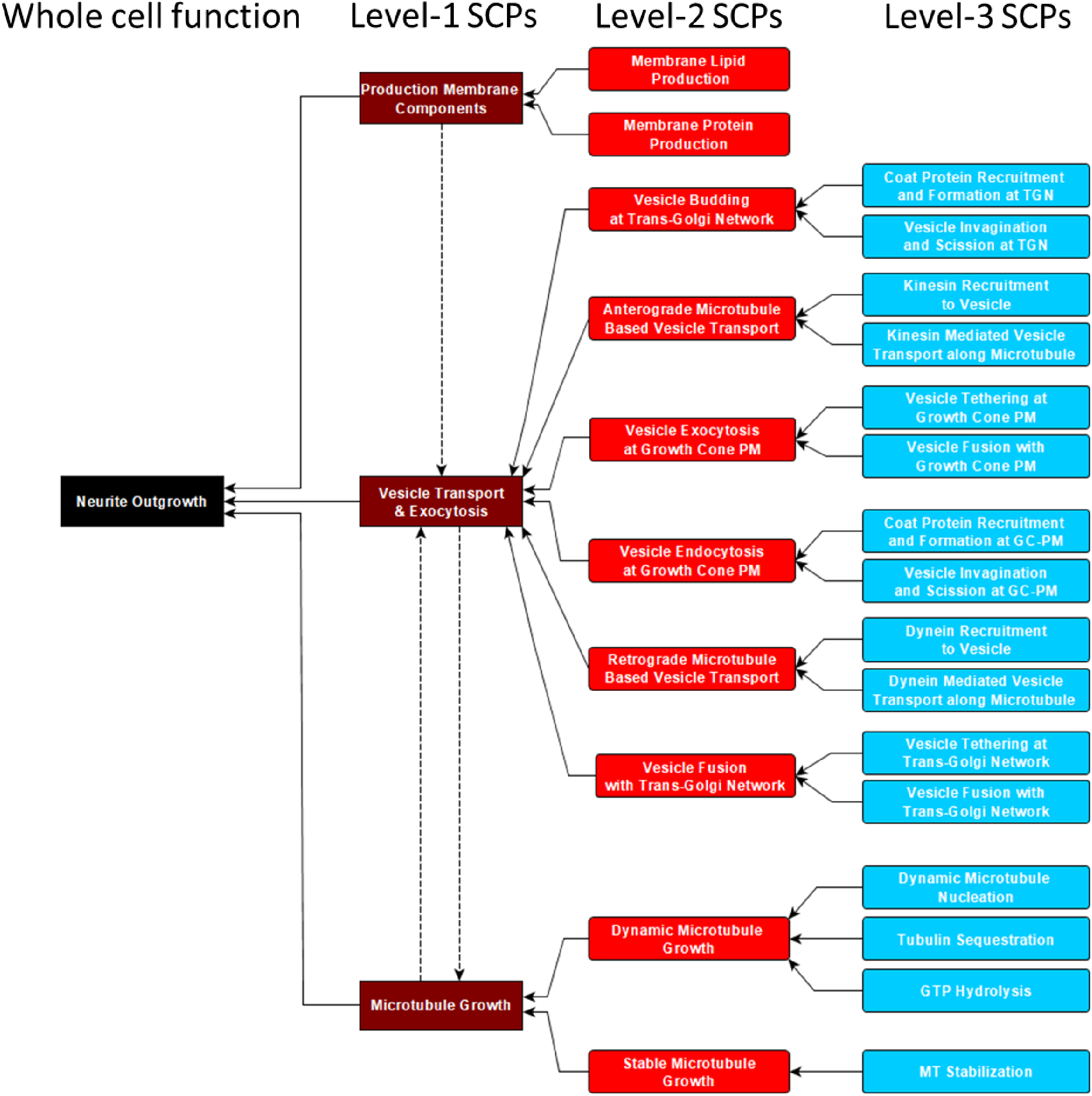
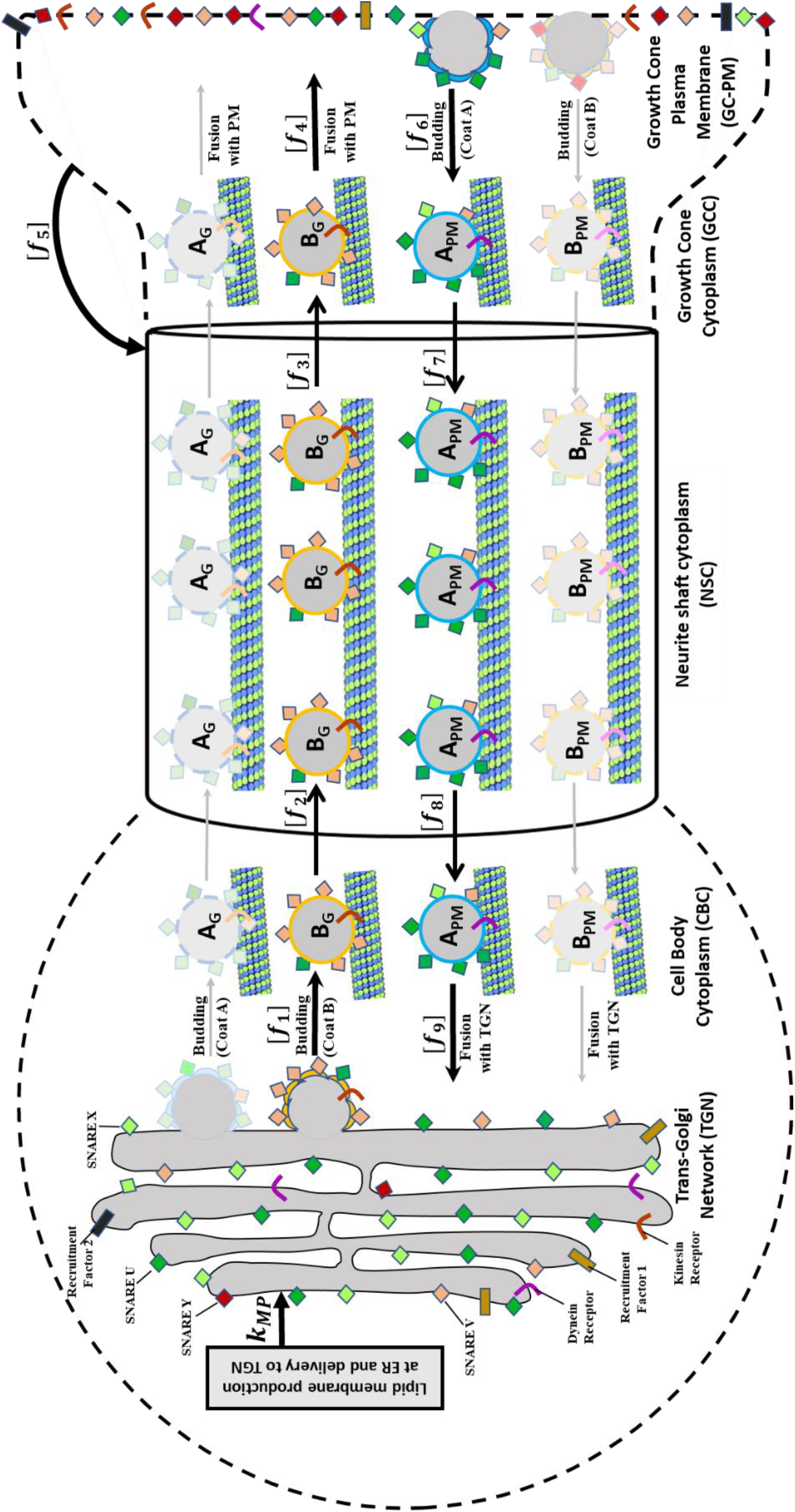
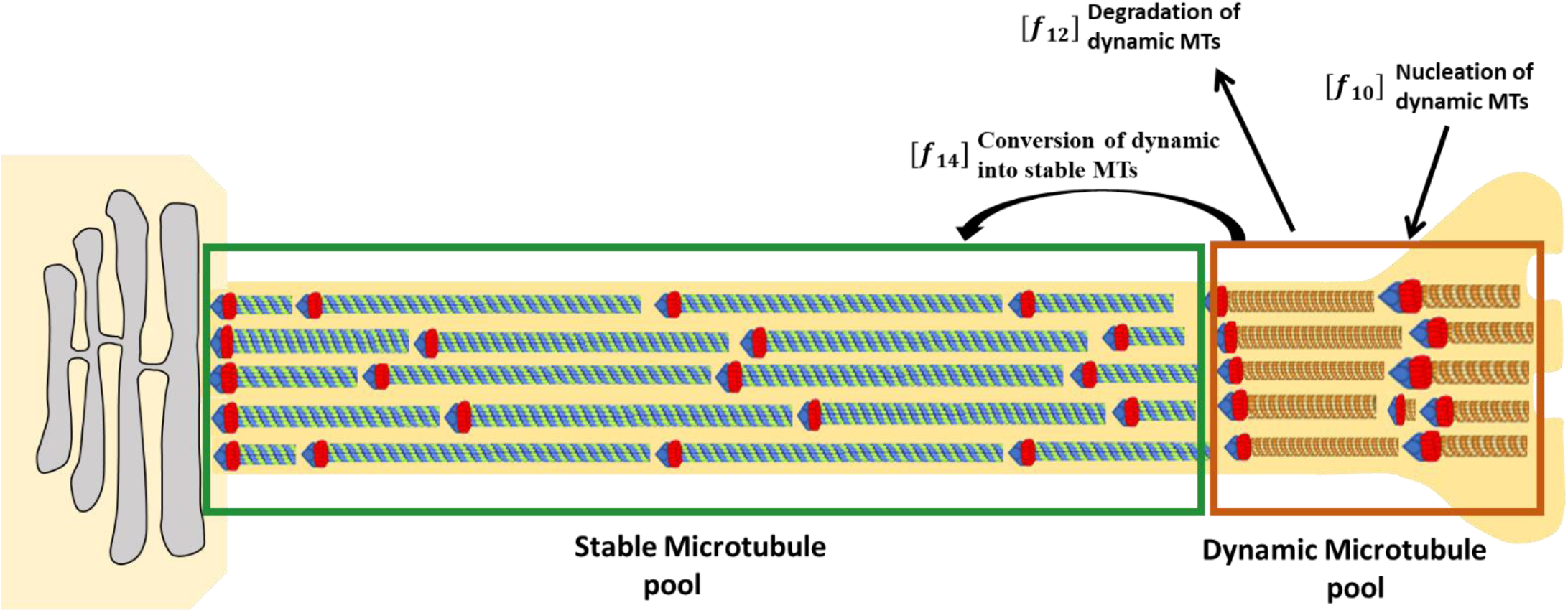
Identification of different level SCPs involved in neurite outgrowth and their incorporation into multicompartmental ODE model. **(A)** Based on our observations dystrophic bulbs (right panel) revealed by tubulin staining are 8–10 times larger than physiological growth cones (left panel). Characteristics of dystrophic bulbs are an accumulation of anterograde vesicles (Bradke et al., 2012) as well as disorganized microtubules (Blanquie and Bradke, 2018)**. (B)** The three major SCP types, *Production of Membrane Components, Vesicle Transport & Exocytosis and Microtubule Growth*, describe essential sub-cellular functions involved in neurite outgrowth. We labeled them as Level-1 SCPs and identified their Level-2 children and Level-3 grandchildren SCPs. **(C)** To model membrane production at the Trans-Golgi network (TGN) and delivery to the growth cone plasma membrane (GC-PM) via vesicle transport, we extended a dynamical model that simulates vesicle transport between the endoplasmic reticulum and the Golgi (Heinrich and Rapoport 2005). In our version, new synthesized membrane is added to the Trans-Golgi network from where it is transported to the growth cone plasma membrane by vesicular transport. Vesicles bud from the TGN, move along the microtubules via active kinesin mediated transport and fuse with the growth cone plasma membrane, leading to neurite shaft growth. The vesicles pass through three intermediate compartments: the cell body cytoplasm (CBC), the neurite shaft cytoplasm (NSC) and the growth cone cytoplasm (GCC). In each compartment kinesin has a different affinity for microtubules, leading to varying fractions of vesicles that are actively transported via kinesin along the MT. Retrograde vesicles are generated via endocytosis at the growth cone plasma membrane and move along the microtubule towards TGN through dynein mediated active transport. Vesicle budding at the TGN or GC-PM is mediated by the interaction of recruitment factors and coat proteins, and vesicle fusion with the TGN or the growth cone membrane by the formation of SNARE complexes between vesicle(v)-SNAREs and target(t)-SNAREs, catalyzed by local tethering machineries. Motor proteins are bound to vesicles through motor protein receptors. A_G_/B_G_ label vesicles that bud from the TGN with coat protein A/B, A_PM_/B_PM_ vesicles that bud from the GC-PM with coat protein A/B. Since at steady state almost all anterograde vesicles bud with coat protein B from the TGN, and almost all retrograde vesicles bud with coat protein A from the GC-PM, we highlighted their transport routes in brighter colors. **(D)** To simulate MT growth, we consider two different MT pools, stable and dynamicMTs. After nucleation new MTs are added to the pool of dynamic MTs that is characterized by alternating phases of growth and catastrophic breakdown. The frequency and duration of these phases depend on the tubulin concentration (regulated by the SCP *Tubulin Sequestration)* and GTP hydrolysis rate (SCP *GTP Hydrolysis)*. Consequently, the length distribution of the dynamic MT pool and the degradation rate of dynamic MTs depends on the activity of both SCPs (supplementary figure 2). Dynamic MTs are either degraded or converted into stable MTs that form the growing neurite scaffold.

The level-1 SCP *Production of Membrane Components* has two Level-2 children SCPs, *Membrane Lipid Production* and *Membrane Protein Production*, that are incorporated into our model via constant synthesis rates of the individual components at the Trans-Golgi Network (TGN) (Figure 1C; Reaction 1 in Table S1) (Gracias et al., 2014; Wang et al., 2011).

The Level-2 children SCPs of the Level-1 SCP *Vesicle Transport & Exocytosis* are *Vesicle Budding at Trans-Golgi Network* and *Vesicle Endocytosis at Growth Cone Plasma Membrane* (GC-PM), *Anterograde* and *Retrograde Microtubule-based Vesicle Transport, Vesicle Fusion with Trans-Golgi Network* and *Vesicle Exocytosis at Growth Cone Plasma Membrane*. Each of these six Level-2 SCPs has two Level-3 children SCPs. To simulate the dynamics of the vesicle transport SCPs we extend a dynamical model of bidirectional membrane lipid and protein transport between the endoplasmic reticulum and the cis-Golgi (Heinrich and Rapoport, 2005). In our model the two organelles are the TGN and the GC-PM (Figure 1C). The level-2 SCP *Vesicle Budding at the TGN* consists of the Level-2 SCPs *Coat Protein Recruitment and Formation at the TGN* and *Vesicle Invagination and Scission at TGN*. The former is mediated by Recruitment Factor 1 that resides at the TGN, the latter is mediated by a cell body specific budding rate (Reaction 2 in Table S1). Membrane proteins at the TGN are recruited into the budding vesicle via association with the coat proteins (Table S2). The Level-2 SCP *Anterograde Microtubule-Based Vesicle Transport & Exocytosis* that moves vesicles from the cell body cytoplasm into the neurite shaft cytoplasm and from there into the growth cone cytoplasm involves the Level-3 SCPs *Kinesin Recruitment to Vesicle* and *Kinesin Mediated Vesicle Transport along Microtubule*. Kinesin motor proteins are recruited to the vesicle via kinesin receptors that are part of the vesicle membrane. Here, we assume that kinesin is available under saturating conditions, so that the kinesin concentration per vesicle is determined by the number of kinesin receptors. Based on the number of kinesin bound kinesin receptors we calculate the likelihood that a vesicle is bound to the microtubule (Table S1, Reactions 6/7). This likelihood also depends on the affinity of kinesin to the microtubule, incorporated into our model as the fraction of Microtubule (MT)-bound kinesin. The fraction of MT-bound kinesin decreases from the cell body cytoplasm to the neurite shaft cytoplasm and growth cone cytoplasm (Table S3). A low fraction of MT-bound kinesin in the neurite shaft cytoplasm ensures that only a few vesicles (~9%) are moving forward, in agreement with experimental data (Ahmed and Saif, 2014). The large pool of stationary vesicles in the neurite shaft cytoplasm could serve as a membrane reservoir that allows quick recruitment of additional membrane to the GC-PM on demand. In the growth cone cytoplasm almost all vesicles are unbound, allowing their fusion with the GC-PM.

The Level-2 SCP *Vesicle exocytosis at GC-PM* is mediated by the interplay of the Level-3 SCPs *Vesicle Tethering* and *Vesicle Fusion at GC-PM*. The former is realized in our model via a growth cone specific tethering rate, the latter via complex formation between vesicle(v)-SNARE V and target(t)-SNARE Y that resides at the GC-PM (Table S1, Reaction 8). Retrograde vesicle transport from the GC-PM to the TGN is incorporated by a similar set of Level-2 and Level-3 SCPs that consist of a symmetric set of membrane proteins and rate constants. We assume a high fraction of dynein is attached to the MT in the neurite shaft cytoplasm, so that most of the retrograde vesicles are actively transported towards the TGN, without formation of a vesicle reservoir.

Vesicles that fuse with the GC-PM, increase its surface area. In our model, we set the GC-PM surface area to 50 μm^2^ (Kunda et al., 2001; Ren and Suter, 2016). Any membrane surface area that is added to the GC-PM and would increase its size beyond 50 μm^2^ is added to the neurite shaft (Table S1, Reaction 9). Membrane proteins are not added to the neurite shaft, since our model is based on the assumption that the diffusion of the membrane proteins into the neurite shaft is prevented by intra-membranous diffusion barriers or cortical cytoskeleton proteins (Ye and Zhang, 2013) that ensure a highly specialized growth cone plasma membrane. Neurite shaft length arises from its membrane surface area (Table S1, Reaction 10). For simplicity, we do not consider increase of the neurite diameter at this stage. The neurite shaft cytoplasm lies within the neurite shaft and grows with the lengthening neurite shaft. Most of the anterograde vesicles in the neurite shaft cytoplasm are not actively transported along the microtubule but are stationary components of the membrane reservoir. Consequently, the growing neurite shaft cytoplasm acts as a sink for vesicles and vesicle membrane proteins.

Microtubule (MT) growth is mediated by the interplay of the Level-2 SCPs *Dynamic Microtubule Growth* and *Stable Microtubule Growth*. (Brady et al., 1984; Kollins et al., 2009; Song et al., 2013; Witte et al., 2008). Dynamic MTs are characterized by periods of MT growth and catastrophic breakdown, the latter can either lead to MT disappearance or rescue, followed by a new growth period. The time frame of these periods is in seconds or minutes. Stable MTs do not show such periodic growth behavior and we assume that there is no degradation of stable MTs. Dynamic MTs are continuously generated at a specified nucleation rate (Level-3 SCP *Dynamic MT Nucleation)* (Reaction 11 in Table S4, Figure 1D). The length of dynamic MTs (Reaction 12, Table S4) as well as the rate of MT catastrophic breakdown depends on the free tubulin concentration that is regulated by the SCP *Tubulin Sequestration* (Reaction 13 Table S4) (Curmi et al., 1997; Jourdain et al., 1997; Manna et al., 2009) and on the hydrolysis rate of tubulin-bound GTP (SCP *GTP Hydrolysis)*. The latter SCP determines the size of the GTP cap at the tip of the dynamic MT that protects it from catastrophic breakdown (Seetapun et al., 2012).

### Conceptual frame work for modeling whole cell dynamics arising from interactions between SCPs

Our model uses whole cell responses in a top-down based manner that started with the literature curation of SCPs involved and their assignment to different levels of biological detail (Figure 2, Step 1a), as described above. Like the organization of our Molecular Biology of the Cell Ontology (Hansen et al., 2017), Level-1 and Level-2 SCPs describe more general, Level-3 SCPs more specific cell biological or biochemical functions. SCPs of the different levels are in parent-child relationships. SCPs are represented as coupled biochemical reactions that are transformed into coupled ordinary differential equations (ODEs) (Figure 2, Steps 1b/c). We use experimental data to define model constraints that the simulations must match (Figure 2, Step 1d). For systematic analyses we develop analytical solutions for the prediction of kinetic parameters of SCPs at steady state (Figure 2, Step 2a). Here, we use the term steady-state to describe a continuous whole cell response without variation of amplitude. We use the analytical solutions to predict steady-state kinetic parameters and values that match the predefined model constraints (Figure 1, Step 2b). Through systematic parameter variation, we generate multiple steady-state solutions that allow for the classification of dependencies between SCPs at different levels (Figure 2, Step 2c). To investigate the effect of SCP dysfunction on whole-cell responses, we generate numerical simulations based on steady-state parameter sets after arbitrary modification of kinetic parameters of a single SCP followed by experimental verification (Figure 2, Step 2d).

**Figure 2:**
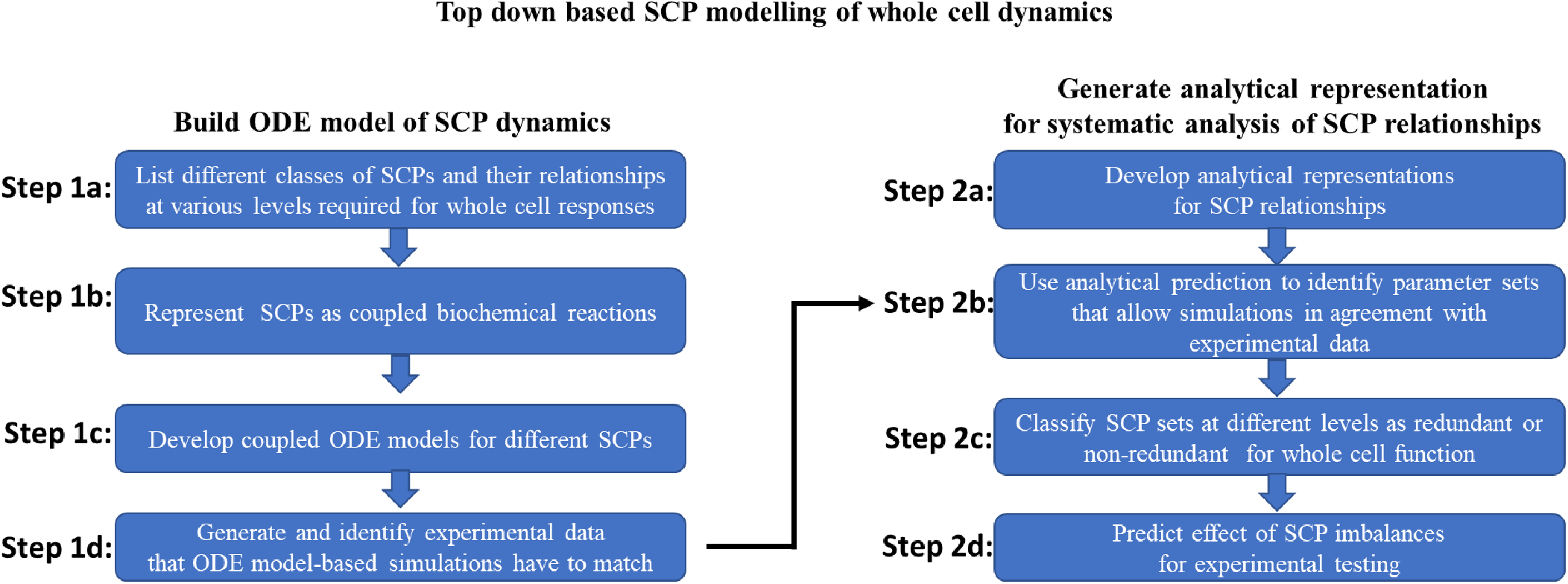
Top-down based SCP modelling of whole cell dynamics. Our pipeline for the generation of dynamical models that allow the analysis of relationships between sub-cellular processes (SCPs) during whole cell responses consists of two major steps. First, we build multicompartmental ODE models for the simulation of SCP activities that are based on literature curated data (Step 1a-d). In the second step we develop analytical predictions for kinetic parameters at steady state that allow a systematic analysis of SCP interactions (Step 2a-d).

### Simulations, Analytical Analyses and Predictions

To characterize the coordinated activities of these Level-3 SCPs, we ran an initial set of simulations of the growth of a single dynamic MT using a stochastic growth model (Margolin et al., 2012) (Figure S1). Since we simulate NOG over hours and days, while dynamic MT growth behavior changes in seconds and minutes, we do not simulate individual dynamic MTs, but the whole population of dynamic MTs. For this population simulation, we developed two formulas based on the results of the model by Margolin et al. They describe the dependence of the average growth length of dynamic MTs and of the catastrophic breakdown rate that leads to complete disappearance of the dynamic MT on the effective tubulin concentration and the GTP hydrolysis rate (Reactions 12/13 in Table S4; Figure S1). The count of dynamic MTs depends on the nucleation rate, the degradation rate due to catastrophic breakdown and the conversion rate into stable MTs (Reaction 14 in Table S4; Figure S1). The activity of the Level-2 SCP *Stable Microtubule Growth* is regulated by the Level-3 SCP *MT Stabilization* (Table S4, Reaction 15). To calculate the length of the microtubule scaffold within the neurite, we sum the length of all dynamic and stable MTs and divide it by the number of MTs per neurite cross-section (Reaction 16/17 in Table S4).

### Experimental determination of varying neurite growth rate velocities

In primary cultures, neurons put out neurites at varying rates. To quantify the variability and determine the velocities with which neurites grow, we used a high content imaging system (IN Cell Analyzer) to track neurite outgrowth in primary rat cortical neurons. The experimentally observed velocities of neurite outgrowth were used as a top-down constraint for our model. We incubated rat cortical neurons for 16h and then documented neurite outgrowth every 6h for an additional 48h. For each neuron, we quantified the length of its longest neurite (the one most likely to become an axon). Representative images are shown in Figure 3A and 3B, and a complete set of images is shown in Supplementary Figure 2. At each time point the neurites significantly differed in their lengths (Figure 3B). We identified the longest neurite for each neuron in a field (Supplementary Figure 2) and calculated their length distribution at every time point (Figure 3C), identified the length of selected quantiles and calculated the growth velocity for these quantiles (Figure 3D). The neurite growth velocities ranged from 0 to ~20 μm/h. We used these velocities as a top-down whole-cell response constraint in modeling the activities of the Level-1, −2 and −3 SCPs. Increase in the whole-cell response, i.e. neurite outgrowth velocity, depends on the coordinated increase of membrane production and delivery of membrane vesicles to the growth cone as well as microtubule growth. Both types of SCPs need to facilitate growth with the same velocity, since they are interdependent. Vesicle transport is microtubule dependent and microtubule growth is limited by the length of the neurite shaft. Similarly, the increase in membrane production needs to be coordinated with increased vesicle transport. More vesicles would need to bud from the TGN, to be actively transported through the neurite shaft cytoplasm and to fuse with the GC-PM. These interdependencies form the basis for our top-down constraint.

**Figure 3:**
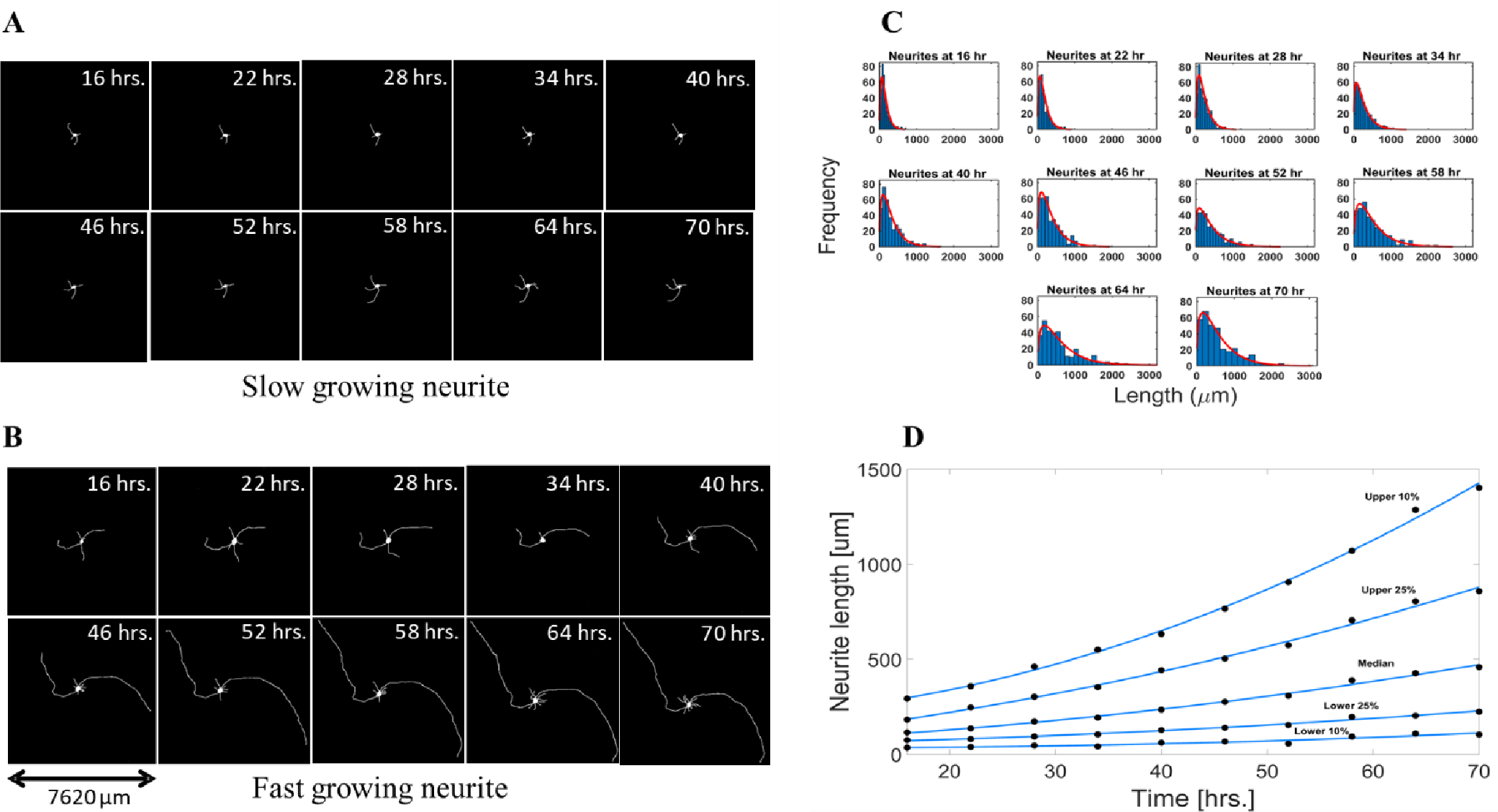
Neurites grow with different outgrowth velocities. Neurons were dissected from rat cortical brain, incubated for 16h to allow initial growth, and followed by image acquisition every 6h up to 70h after plating. **(A-B)** The growth dynamics of two example neurons, one with a low (A) and one with a high (B) outgrowth velocity, are shown over given time-period. **(C)** At each timepoint we quantified the length of the longest neurite of each neuron. Histograms show the length distribution of all longest neurites at the indicated timepoints. **(D)** At each timepoint we identified the top and bottom 10% and 25% quantile length as well as the median length. Linear interpolation of the lengths for each selected quantile documented outgrowth velocities ranging from 0 to ~20 μm/h.

### Postulates governing SCP dynamics

In addition to the experimentally determined NOG velocities, we used experimental data from the literature for additional constraints at the level of SCPs. These constraints arise from a set of postulates of permissible SCP functions that are described in (Table S5). We postulate that about ~10% of the anterograde vesicles in the neurite shaft cytoplasm are bound to the microtubule and actively moving along the microtubule (Ahmed and Saif, 2014). The other 90% constitute a membrane pool that can be recruited on demand to adapt to short-term increases in outgrowth velocity. Similarly, we assume that the vesicles that reside in the growth cone (Erturk et al., 2007) are the first membrane reservoir for the quick recruitment of additional membrane and also define - in the same way - that 10% of the anterograde vesicles in the growth cone cytoplasm move, i.e. fuse with the growth cone membrane. This assumption is based on the experimental observation that GC membrane precursor vesicles reside in the GC cytoplasm before fusion with the membrane with a half-life of ≥14min (Pfenninger et al., 2003), which would suggest that around 5% of the vesicles fuse with the membrane. Similarly, we assume that 90% of the retrograde vesicles in the neurite shaft cytoplasm and the cell body cytoplasm are moving. The total growth cone membrane is completely internalized within 30 - 60 min (Diefenbach et al., 1999) which suggest an endocytosis rate between 0.833 μm^2^/min - 1.66 μm^2^/min for a growth cone with a surface area of 50 μm^2^ (Kunda et al., 2001; Ren and Suter, 2016). Based on these observations and the assumption that some of the endocytosed vesicles might be part of transcytotic re-arrangements during growth cone steering (Tojima and Kamiguchi, 2015) or are transported to other growth cones and not to the cell body (Denburg et al., 2005), we set the rate of membrane back transport from the growth cone to the TGN to 0.5 μm^2^/min. The backward transported membrane can also be called cycling membrane, since it continuously cycles between the TGN and the GC-PM without being added to the neurite shaft. For the microtubules we postulate that dynamic MTs should fill up 20 μm of the growing neurite. This is the length of the minor processes (Yu and Baas, 1994) that are characterized by continuous retraction and extension periods (Arimura and Kaibuchi, 2007). We assume that such periods are committed by continuous MT growth and retraction, so that the MT scaffold in these processes should mainly consist of dynamic MTs.

Using these constraints, we ran simulations to identify an initial solution at one intermediate rate of neurite outgrowth. From these initial simulations, we developed an overall analytical approach that allows us to predict the relationships that produce neurite outgrowth at a specified velocity.

We identified a set of parameters that generated neurite outgrowth with a velocity of 10 μm/h (Figure 4 and S3). Under steady-state conditions neurite, surface area continuously increased at a rate of 0.5236 μm^2^/min in agreement with the amount of membrane that is needed to increase the length of the growing neurite (Figure 4A). The length of the microtubule bundles also increase proportionally to maintain the shaft growth rate (Figure 4B). For this we selected an effective tubulin concentration of 9 μM to achieve a length increase of the microtubule bundles with a velocity of 10 μm/h. The vesicles in the neurite shaft cytoplasm increased in parallel with a rate of ~0.57 vesicles/min assuming a membrane area of 0.05 μm^2^ per vesicle (Figure 4C). In considering the values of the ordinate for the different SCPs (Figure 4) it became clear that there are multiple combinations of kinetic parameters and amounts of components that allow neurite outgrowth at the same velocity without violating the constraints we imposed on dynamics of the SCPs.

**Figure 4:**
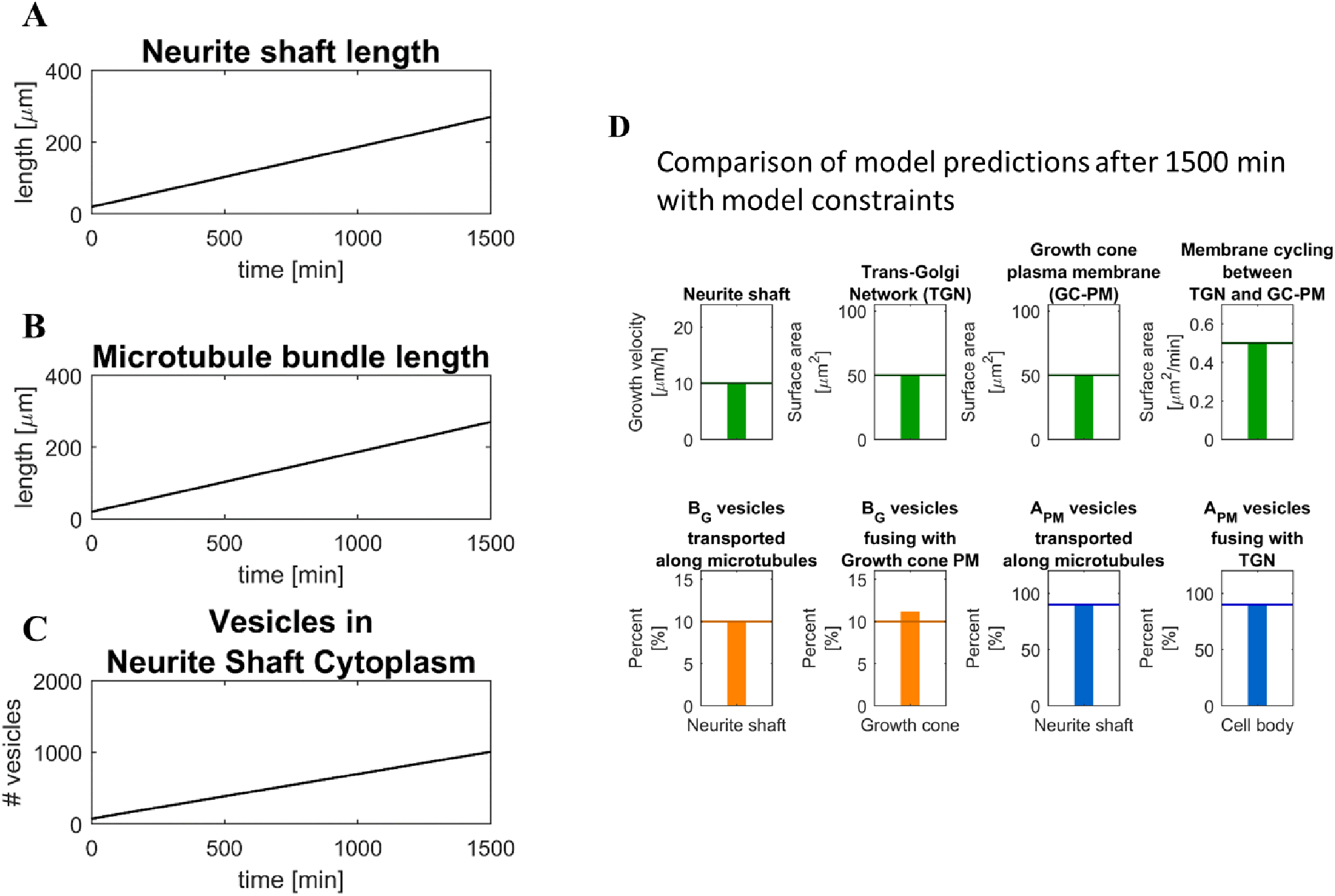
Simulation of vesicle transport and microtubule growth during neurite outgrowth. **(A-B)** We identified a set of kinetic parameters (Supplementary table S7) that allows neurite outgrowth via coordinated neurite shaft (A) and microtubule bundle growth (B) with a velocity of 10 μm/h. **(C)** The growing neurite shaft cytoplasm (NSC) within the growing neurite acts as a sink for anterograde and retrograde vesicles, causing an accumulation of vesicles within this compartment. **(D)** Our identified set of kinetic parameters allows NOG without violation of the model constraints that we selected to define physiological outgrowth conditions.

To enable a systematic analysis of the redundancies within and between SCPs, we generated an analytical solution (Figure 5, and S4) for the prediction of kinetic parameters at steady state that enable neurite outgrowth at a given velocity without violating constraints on SCP dynamics (Table S5). In our solution, we consider that an increase in neurite outgrowth depends on varying sets of protein amounts and kinetic parameters that allow increased net anterograde transport of membrane vesicles and continuous synthesis of new proteins and membrane to compensate for the accumulation within the growing neurite shaft cytoplasm. In addition to the postulated constraints on SCP dynamics, we pre-define independent protein amounts and kinetic parameters (Figure S4, right side) to calculate the dependent protein amounts and kinetic parameters. To achieve a given NOG velocity a certain amount of lipid membrane needs to be synthesized and transported from the TGN to the GC. In addition to this membrane that will be added to the neurite shaft three other membrane types can be classified based on their transport destination, allowing us to distinguish four membrane types (Figure 5A). These membrane types should not be confused with the four-different vesicle sets that transport the membrane between the TGN and GC-PM. Type I membrane buds from the TGN and is added to the vesicle reservoir in the growing neurite shaft cytoplasm. Type II membrane is transported from the TGN to the GC-PM and added to the neurite shaft as anterograde vesicles. Type III membrane is transported anterogradely from the TGN to the GC-PM, endocytosed at GC-PM and added to the NSC as retrograde vesicles. Type IV membrane is transported from TGN to GC-PM and from there back to TGN, and therefore is labeled cycling membrane. Initial and final anterograde and retrograde rates that describe budding and fusion processes at the TGN and GC-PM can be calculated by summing up the involved membrane types. The fluxes of the individual membrane types can be calculated based on the anticipated NOG velocity and model constraints (Figure 5B, (1)). To allow the fusion of sufficient anterograde vesicles with the GC-PM, a certain number of SNARE complexes need to be formed between v-SNAREs V and t-SNAREs Y (Figure 5B, (2)). Consideration of the tethering rate allows the calculation of the needed amount and concentration of v-SNARE Vs. The concentration allows the determination of the initial and final forward fluxes of SNARE V ((Figure 5B, (3)). Similarly, concentrations and fluxes can be calculated for the v-SNARE U that mediates vesicle fusion at the TGN ((Figure 5B, (4)/ (5)). During vesicle budding at the TGN and GC-PM, SNAREs compete with each other for the vesicle binding spots. Knowledge of the fluxes of both v-SNAREs allows the calculation of their concentrations at the TGN and GC-PM (Figure 5, (6)/ (7)). The production rates of v-SNARES (and any other membrane vesicle membrane proteins) can be calculated based on their vesicle concentrations and the number of vesicles that accumulate in the growing neurite shaft cytoplasm (Figure 5B, (8)).

**Figure 5:**
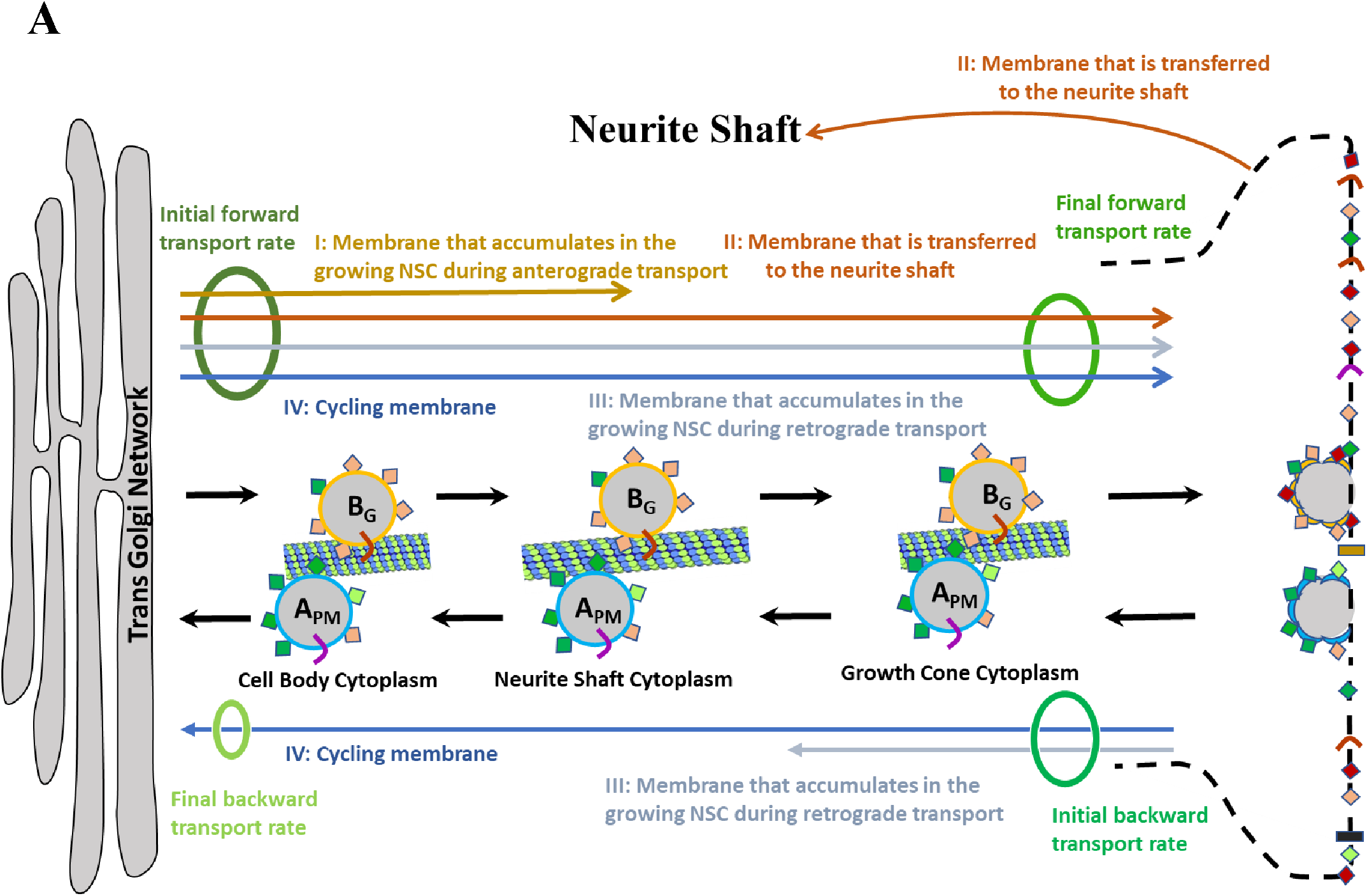
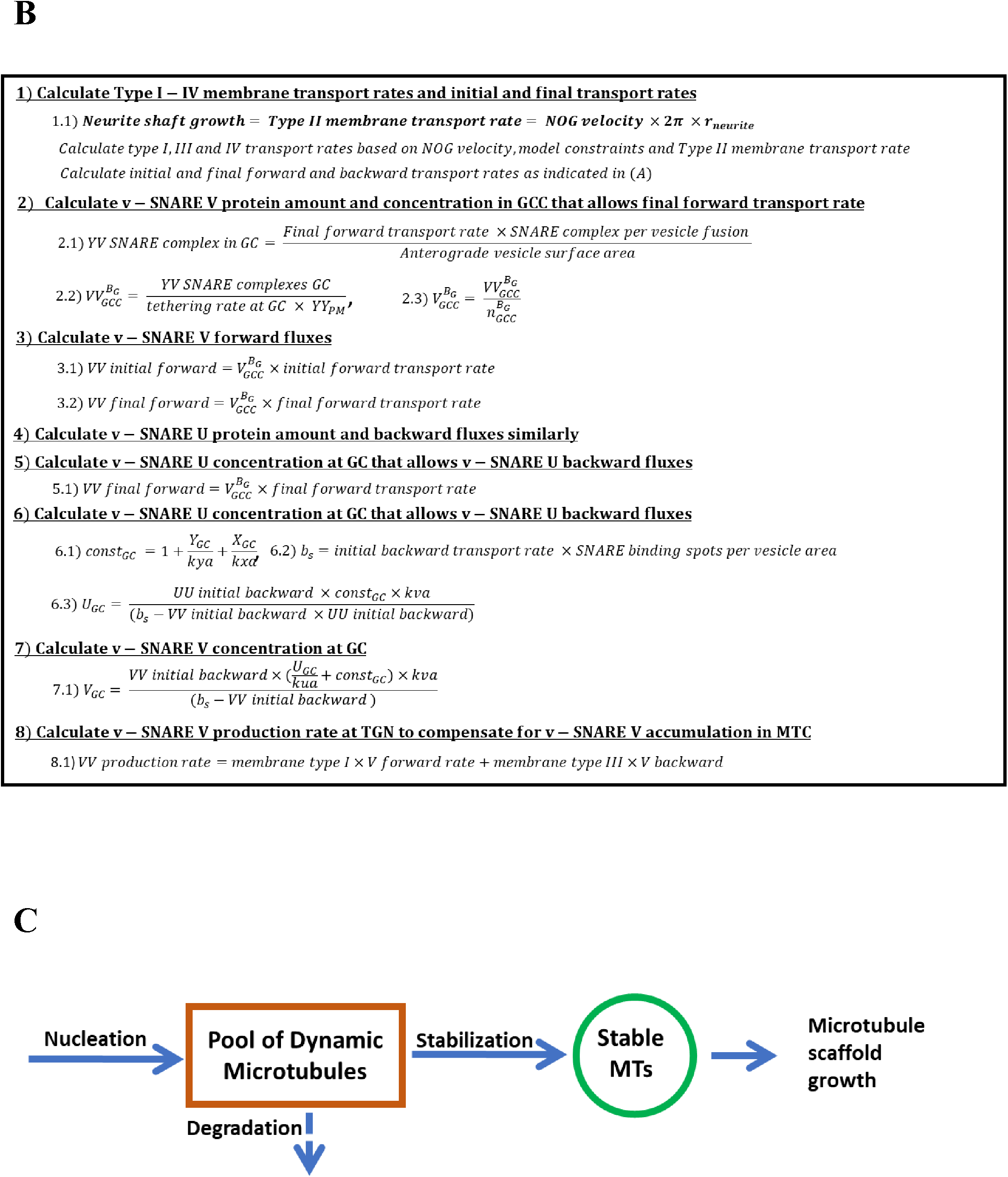
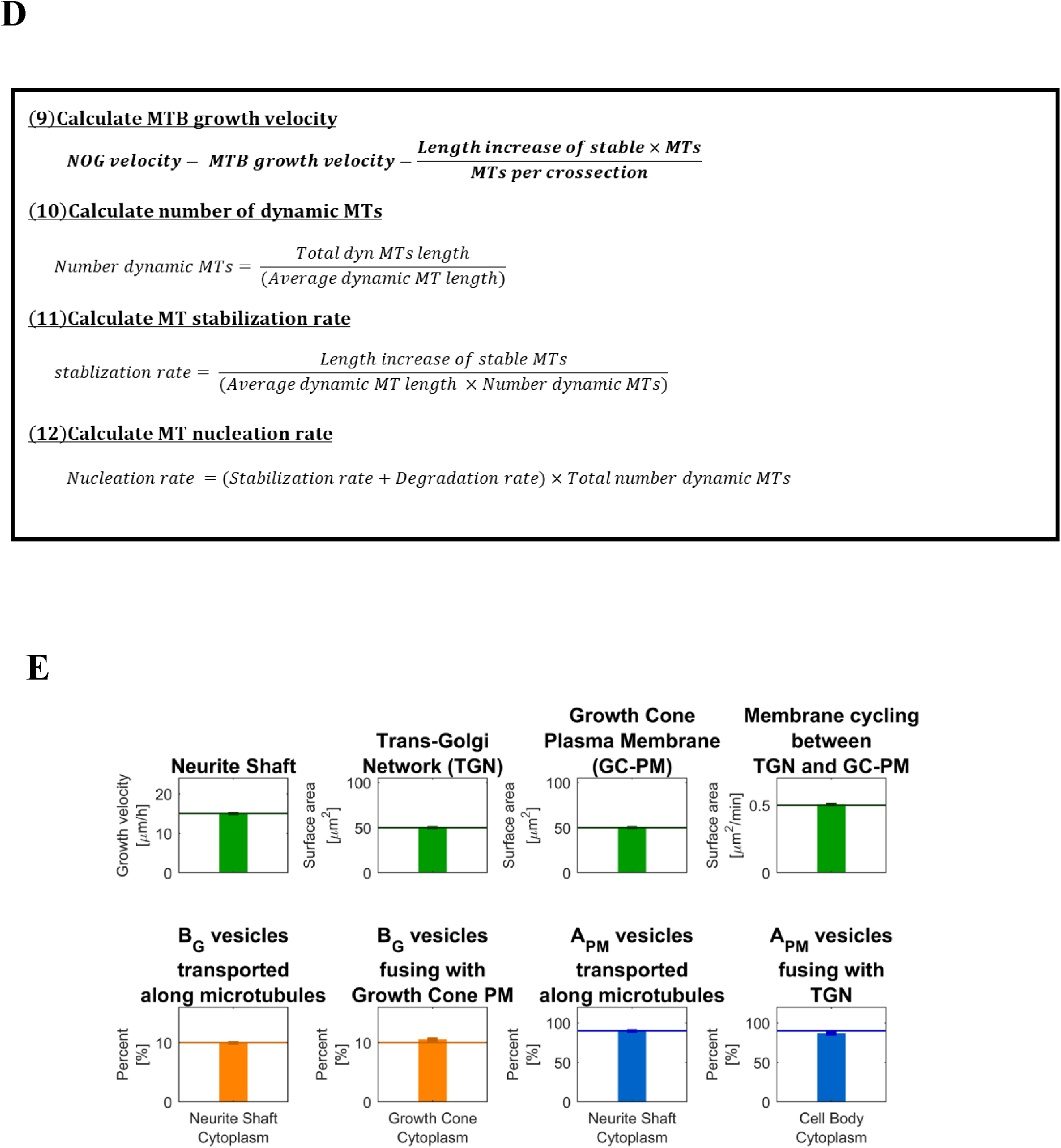
Generation of an analytical solution for the prediction of kinetic parameters at steady state. **(A)** The development of the analytical solution depends on the categorization of membrane fluxes based on their destination compartment. Such classification distinguishes four different membrane types (I-IV) (that should not be confused with the four different vesicle types we show in figure 2B). Initial and final anterograde and retrograde transport rates (which refer to budding and fusion rates at the TGN and GC) are the sum of the transport rates of the encircled membranes types. **(B)** Calculation of kinetic parameter for v-SNARE V. VV/YY/UU refer to total count of SNAREs, V/Y/U describe concentration of SNAREs (per membrane area). See supplementary methods for complete derivation of analytical solution. **(C)** Microtubule bundle (MTB) growth is mediated by the conversion of dynamic MTs into stable MTs. Dynamic MTs nucleate de-novo, followed by their degradation or stabilization. **(D)** Nucleation, stabilization and degradation rates for dynamic MTs are calculated as shown. **(E)** To verify our analytical solution we analyzed, whether our predicted parameter sets allow neurite outgrowth with the selected velocity and without violation of model constraints. For each velocity, we selected 6 different parameter sets that differ in the number of v-SNARE V and the tethering rate at the growth cone plasma membrane (supplementary figure 5A). Every 500 min over a simulation period of 5,000 min (~3.5days), we documented the indicated outputs. Averages and standard deviations were calculated for each velocity. As shown for a velocity of 15 μm/h, our analytical solution predicts steady state dynamics that match the anticipated model constraints (horizontal lines) with high accuracy.

MT growth at a given velocity (Figure 5C) depends on the conversion rate of dynamic into stable MTs (Figure 5D, (9)). By pre-simulation results using the model of Margolin et al, we obtained the dependence of the average length of dynamic MTs on the free (i.e. not sequestered) tubulin concentration and GTP hydrolysis rate. The average length allows the calculation of the number of dynamic MTs (Figure 5D, (10)), followed by calculation of the stabilization rate (Figure 5D, (11)) and subsequently the nucleation rate (Figure 5D, (12)).

The analytical solution was validated by comparing the analytically predicted outcome with the results of the numerical simulation for example sets of parameter and protein amounts (Figure 5E and S5). We used the predicted parameter and protein amount sets as starting points in our model and ran the numerical simulation for 5000 minutes (~3.5 days). Every 500 min we compared the anticipated model constraints with the numerically obtained values to estimate the accuracy of our analytical prediction. The analytical solution matched the anticipated model constraints with high accuracy, except the fusion rate for retrograde vesicles with the TGN that for was too high for the velocities 0 and 2.5 μm/h.

### Complementary Compensatory Relationships between Level-3 sibling SCPs can contribute to robustness of response

We used the analytical solution to systematically analyze the relationship between different SCPs (Figure 1B). For our studies we selected growth velocities that covered the whole range of experimentally observed velocities (Fig 6). We predicted that Level-3 sibling SCPs would have complementary activities. An increase in the activity of one Level-3 SCP can be compensated by a decrease in the activity of at least one of its sibling Level-3 SCPs. To acquire a specified NOG velocity without violation of the model constraints there are multiple combinations of the activities of the two Level-3 SCPs *Coat Recruitment and Formation* and *Vesicle Invagination and Scission* (Figure 6A), the two children SCPs of the Level-2 SCP *Vesicle budding at the TGN*. Such SCP redundancies are also observed between the Level-3 SCPs *Kinesin Recruitment to the Vesicle* and *Kinesin Mediated Vesicle Transport along the MT* (Figure 6B) that define the activity of the Level-2 SCP *Anterograde Microtubule-Based Vesicle Transport*. Similarly, the SCPs *Vesicle Tethering* and *Vesicle Fusion* (Figure 6C), sub processes of the SCP *Vesicle Exocytosis* are characterized by an interaction that is based on an inverse relationship between their activities.

**Figure 6:**
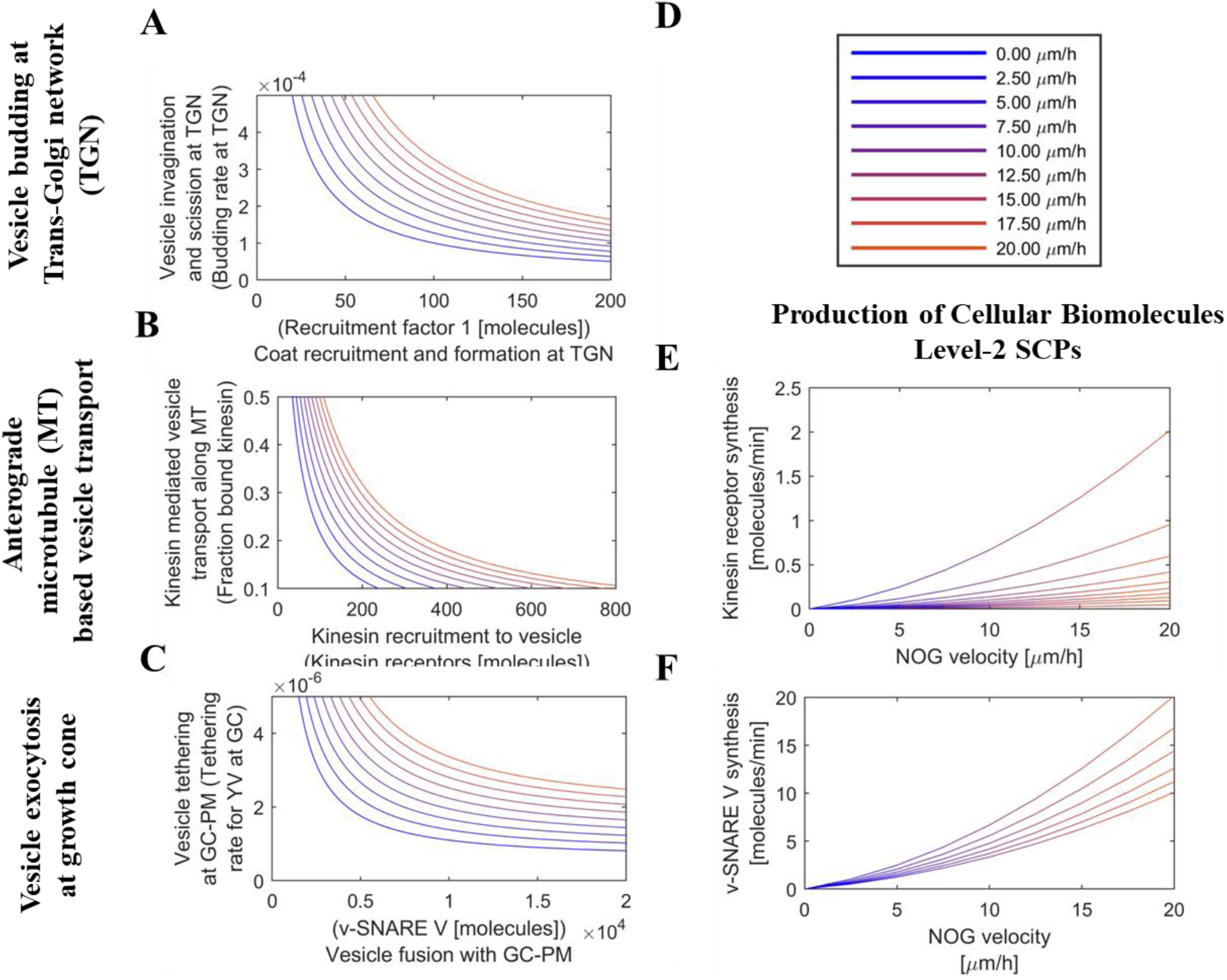
Redundancies among vesicle transport level-3 sibling SCPs. To analyze the relationships between Level-3 SCPs, we used our analytical solution to predict steady state dynamics that enable neurite outgrowth for specified velocities. **(A), (B) & (C)** Level-3 SCPs show complementary relationships between their activities to generate higher level SCP function without violation of the model constraints. **(A)** Different activities of the SCPs *Coat Recruitment and Formation at TGN* and *Vesicle Invagination and Scission at TGN* were modeled by changing the amount of Recruitment Factor 1 and the budding rate at the TGN. SCP activities were inversely related and increased with increasing velocity. Line colors indicate selected velocities as shown in (G). **(B)** A similar SCP redundancy was observed for the SCPs *Kinesin Recruitment to Vesicle* (simulated by # number of kinesin receptors) and *Kinesin Mediated Vesicle Transport along the MT* (simulated by fraction of bound kinesin). **(C)** Inverse relationship between SCP activities was also observed for the SCPs *Vesicle Tethering* (simulated via the tethering rate at the GC) and *Vesicle Fusion* (simulated by amount # of v-SNARE V). **(D)** Velocities were color coded in each figure as indicated. **(E) & (F)** The more a Level-2 SCP activity is generated by a Level-3 SCP that contains a vesicle membrane protein, the higher the SCP activity of membrane protein production to compensate for the loss of membrane protein in the growing NSC reservoir. **(E)** In dependence of the selected fraction of MT-bound kinesin, vesicles need a different amount of kinesin receptors to ensure NOG without violation of the model constraints. A higher kinesin receptor concentration per vesicle directly translates into the need for a higher kinesin production rate. Lines refer to predefined fractions of MT-bound kinesin: 0.1, 0.2, 0.3, 0.4, 0.5, 0.6, 0.7, 0.8, 0.9, 1.0 (from top to bottom). **(F)** Different tethering rates demand different v-SNARE V concentrations per vesicle and consequently different V production rates to ensure NOG without violation of the model constraints (t-SNARE Y is kept constant). Lines refer to different tethering rates: 2.5 ×10^−6^, 3 ×10^−6^, 3.5 ×10^−6^, 4 ×10^−6^, 4.5 ×10^−6^, 5 ×10^−6^ (from top to bottom).

The balance of the Level-3 SCP activities to fulfill the Level-2 SCP function is not isolated at Level-3 but influences the activity of other SCPs as well. The more the activity of a Level-2 SCP arises from a Level-3 SCP that involves vesicle proteins, the higher the need for continuous membrane protein supply to compensate for the consumption of membrane proteins in the neurite shaft cytoplasm sink. This accounts for the kinesin receptor (Figure 6E) as well as for the v-SNARE V (Figure 6F). Here, one might consider that the complementary SCP *Kinesin-Mediated Vesicle Transport along the MT* is determined by components along the whole microtubule while the complementary SCP *Vesicle Tethering* at GC-PM is determined by components of the tethering machinery that are locally restricted to the growth cone. The maintenance of the activity of the former SCP might therefore depend on a continuous synthesis of cytoplasmic protein that is not considered in our model.

Similar compensatory relationships exist between Level-3 sibling SCPs involved in Microtubule Growth. The combined length of dynamic MTs that are available for conversion into stable MTs is regulated by an inverse relation between the activities of the Level-3 SCPs *Dynamic MT Nucleation*, *GTP Hydrolysis* and *Tubulin Sequestration*. The average dynamic MT length (Figure 7B) as well as the degradation rate of dynamic MTs (Figure 7C) depends upon the Level-3 SCPs *Tubulin Sequestration* (which regulates effective tubulin concentration) and *GTP Hydrolysis*. The balanced activities of these two Level-3 SCPs and the Level-3 SCP *Dynamic Microtubule Nucleation* (Figure 7D/E) determines the activity of the Level-2 SCP *Dynamic Microtubule Growth*. Similarly, to the vesicle transport Level-3 SCPs, any loss of function in one of these MT Level-3 SCPs can be compensated by a gain of function in one of the other.

**Figure 7:**
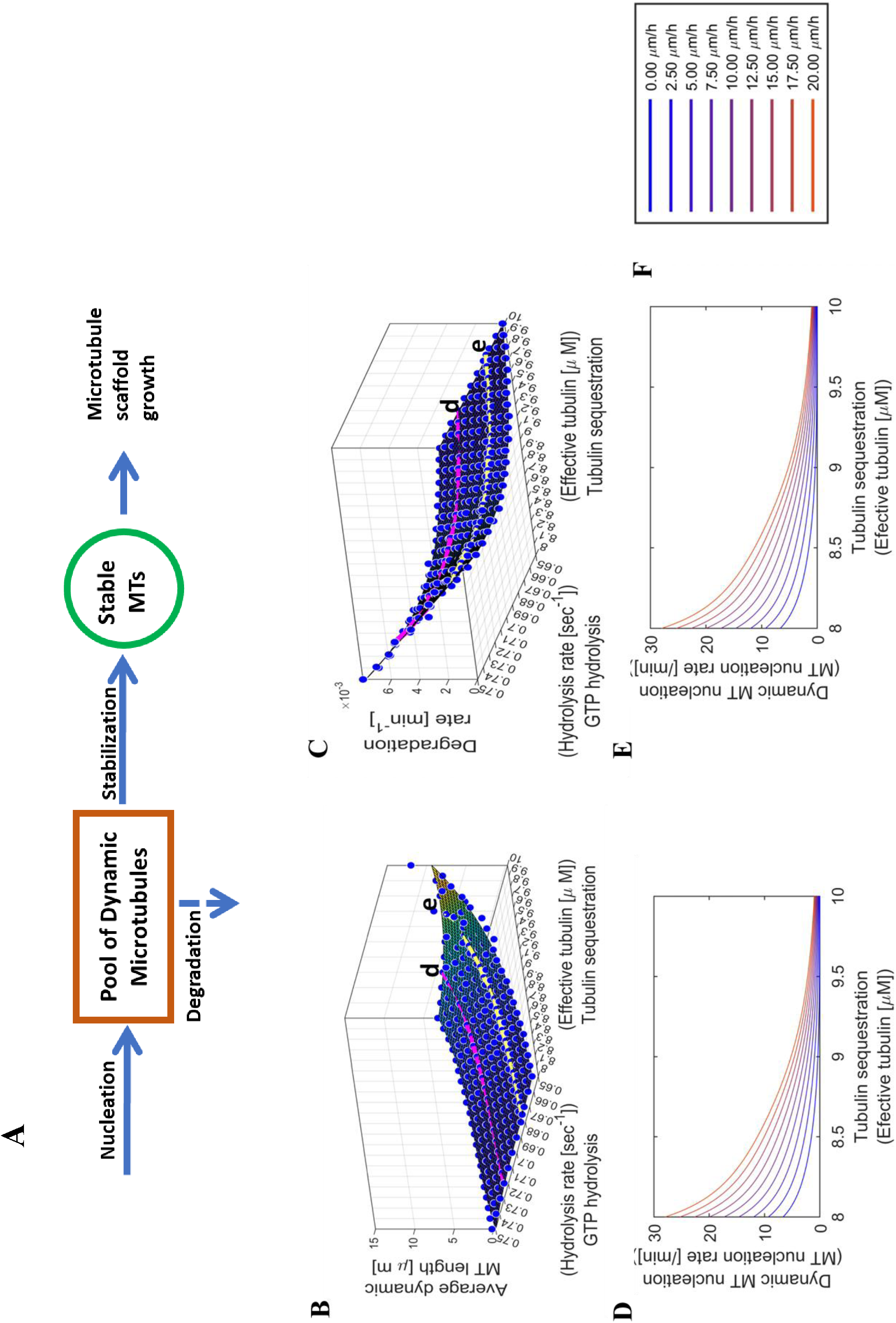
Redundancies among microtubule growth Level-3 sibling SCPs. **(A)** Summary of the basic reaction for the simulation of MT growth. See Figure 2C for details. **(B), (C), (D) & (E)** Level-3 SCPs that define sub-functions of the Level-2 SCP *Dynamic MT growth* show complementary activities to allow NOG without violation of the model constraints. Multiple combinations of the SCP activities *Tubulin Sequestration, GTP Hydrolysis* and *Nucleation of Dynamic MTs* enable NOG with a given velocity. **(B-C)** The influence of the SCPs *Tubulin Sequestration* (which determines the effective tubulin concentration) and *GTP Hydrolysis* on the length distribution (B) and degradation rate (C) of the dynamic MTs were determined in a presimulation using a stochastic model for the simulation of single dynamic MT growth (Margolin et al). Colored lines refer to the selected GTP hydrolysis rates shown in (B) and (C) (pink: 0.72/sec, d) and (yellow: 0.675/sec, e). **(D-E)** Multiple combinations of dynamic MT nucleation rates and effective tubulin concentrations allow NOG under a given velocity for each GTP hydrolysis rate, as shown for the hydrolysis rate of 0.72/sec (d) and 0.675/sec (e). **(F)** Velocities in (D) and (E) were color coded as indicated.

### Sibling Level 2 SCPS do not readily complement each other leading to dystrophic bulbs

The relationships between Level-2 SCP activities are more complicated. A decrease in the activity of one Level-2 SCP cannot simply be compensated by increase in the activity of one of its Level-2 sibling SCPs. To simulate mismatching activity of a Level-2 SCP on NOG, we first predicted a set of kinetic parameters that allow growth under physiological conditions as defined by our model constraints. Then we selected one of its Level-3 SCP children, followed by the perturbation of one kinetic parameter that belonged to this Level-3 SCP, so that this parameter no longer matched anymore to the predicted parameter set. The resulting unbalanced activities between the redundant Level-3 SCP siblings propagated to impaired Level-2 SCP activity that did not match with the activities of the other Level-2 SCPs. We then searched for adaptations in other parameters that allow restoration of neurite outgrowth under physiological conditions as defined by our model constraints. As shown for the example Level-2 SCP *Vesicle Budding at the Trans-Golgi Network*, multiple Level-2 SCP activities needed to be altered in a coordinated manner (Figure 8), e.g. *Membrane Lipid* and *Protein Production* (simulated via reduced production rates), *Anterograde Vesicle Transport along the Microtubule* (simulated via reduced amount of Kinesin receptor) and *Vesicle Exocytosis at the GC-PM* (simulated via reduced amount of v-SNARE V). These activity changes were associated with a reduction of NOG velocity (from 10 μm/h to 5.5 μm/h), since the cycling rate was kept constant at 0.5 μm^2^/min.

**Figure 8:**
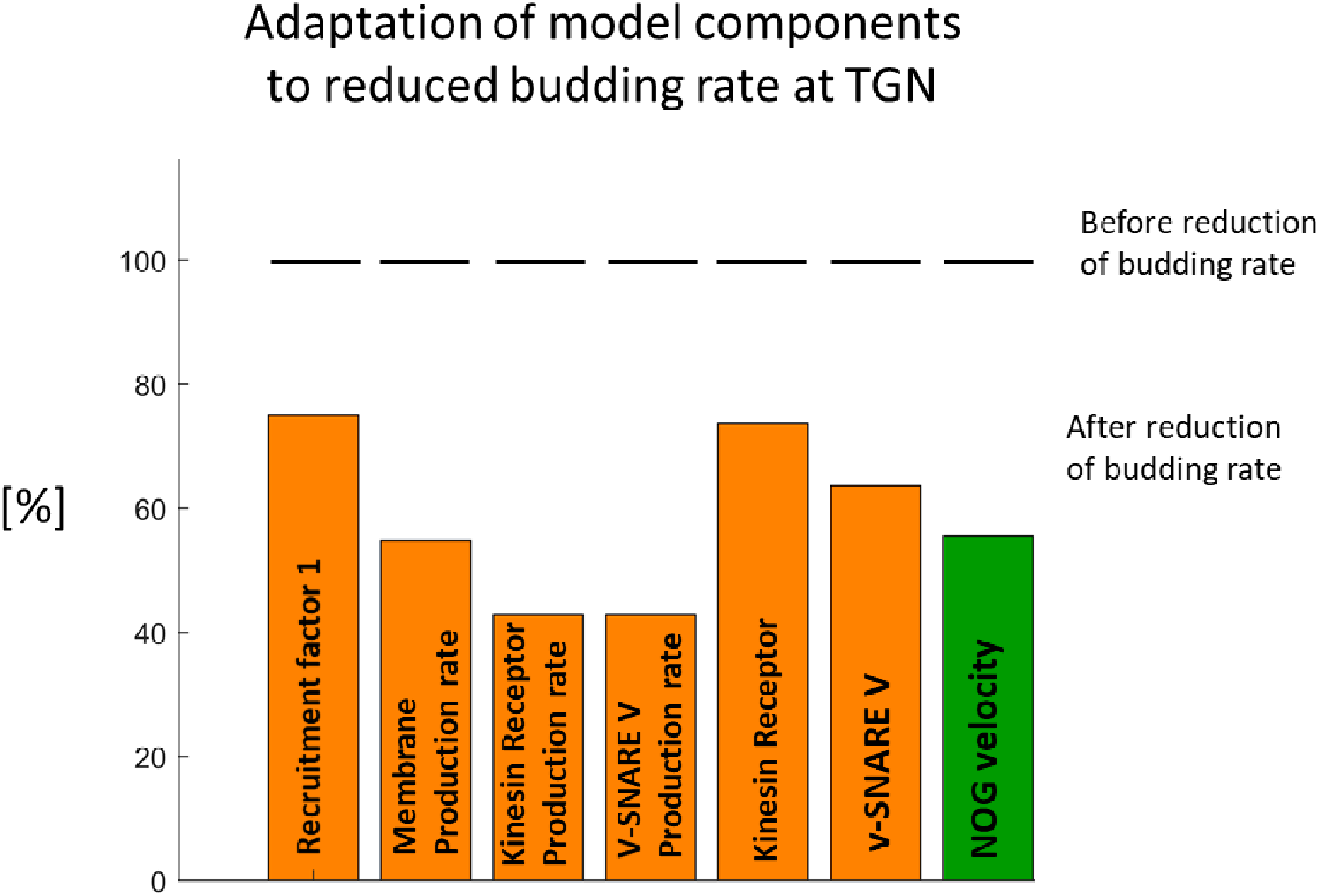
Loss of function of Level-2 SCPs requires multiple adaptations to allow NOG under physiological conditions. We predicted a parameter set that allows steady neurite outgrowth with a velocity of 10 μm/h. To investigate Level-2 SCP dependencies we reduced the activity of the SCP Vesicle budding at the TGN by reducing the amount of Recruitment Factor 1 without compensatory increase of the budding rate. In consequence, the activities of multiple Level-2 SCPs needed to be coordinatively adapted to secure neurite outgrowth under physiological conditions (as defined by our model constraints). One possible adaptation involves the reduction in the activity of the SCPs *Membrane Lipid Production*, of *Anterograde Microtubule-Based Vesicle Transport* (via reduced kinesin receptors), of *Vesicle Exocytosis* (via reduced v-SNARE V) and of *Membrane Protein Production* of the involved transmembrane molecules. Since this option includes a constant cycling rate (i.e. rate of back transported membrane), it causes a reduction in neurite outgrowth velocity.

To simulate the effect of mismatching Level-2 SCP activities on NOG, we reduced the activity of one Level-2 SCP (as described above, but without adaptation of any other Level-2 SCP activities). We selected one *Vesicle Transport & Exocytosis* SCP and one *Microtubule Growth* SCP for further investigation. To simulate the effect of the mismatching activity of the Level-2 SCP *Vesicle Exocytosis* at GC-PM, we selected the v-SNARE V that is involved in the activity of its Level-2 child SCP *Vesicle Fusion at the Growth Cone*. We predicted that a reduction in its activity does not influence NOG velocity, but overloads the growth cone with anterograde vesicles, a feature associated with dystrophic bulbs (Bradke et al., 2012) (Figure 9A/B). To test our predictions, we selected the v-SNARE VAMP7 that is involved in vesicle fusion at the GC-PM (Burgo et al., 2012). In agreement with our predictions, VAMP7 knock down (KD) did not significantly decrease NOG velocity, as shown by the total length of neurites 48h after axotonomy (Figure 9C/D). Its main effect was the generation of dystrophic bulbs (Figure 9C/E).

**Figure 9:**
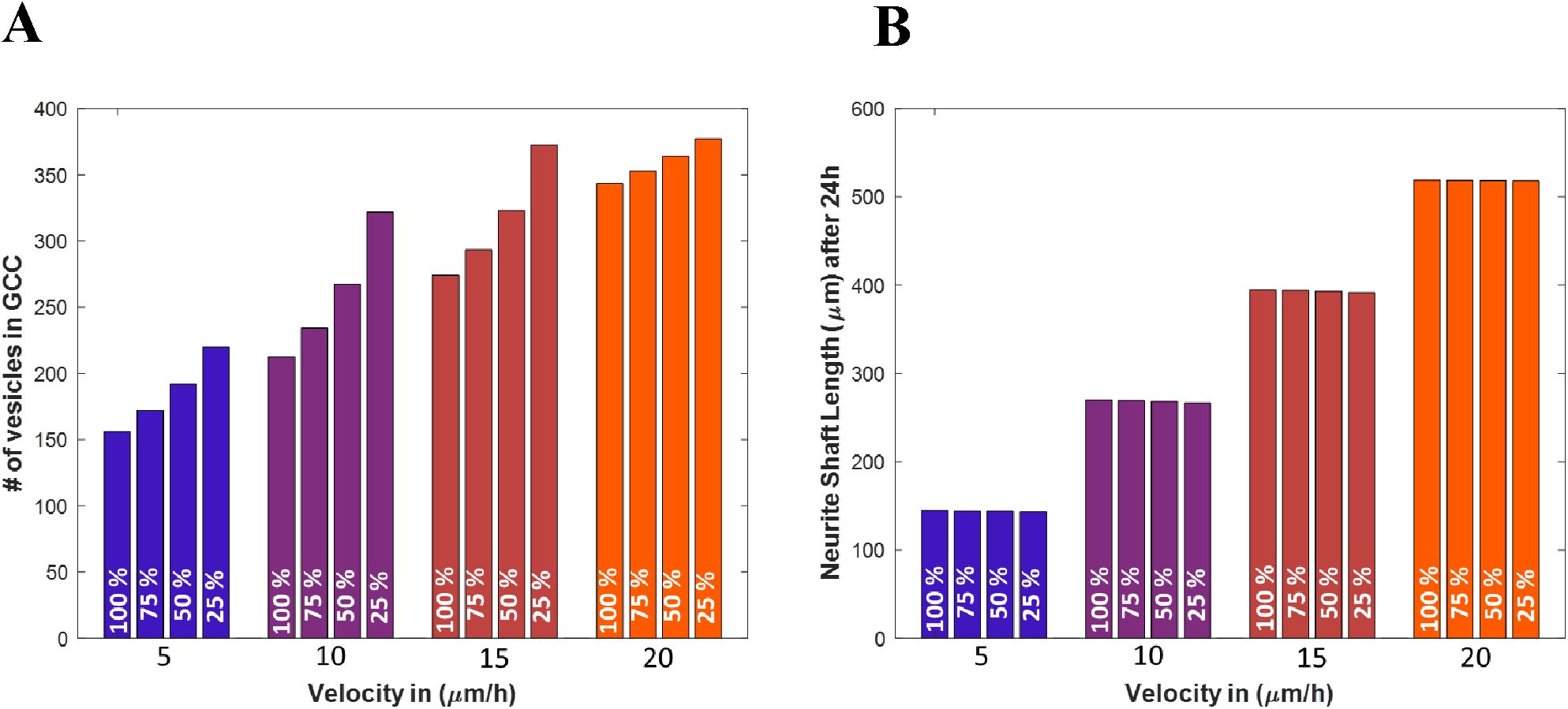
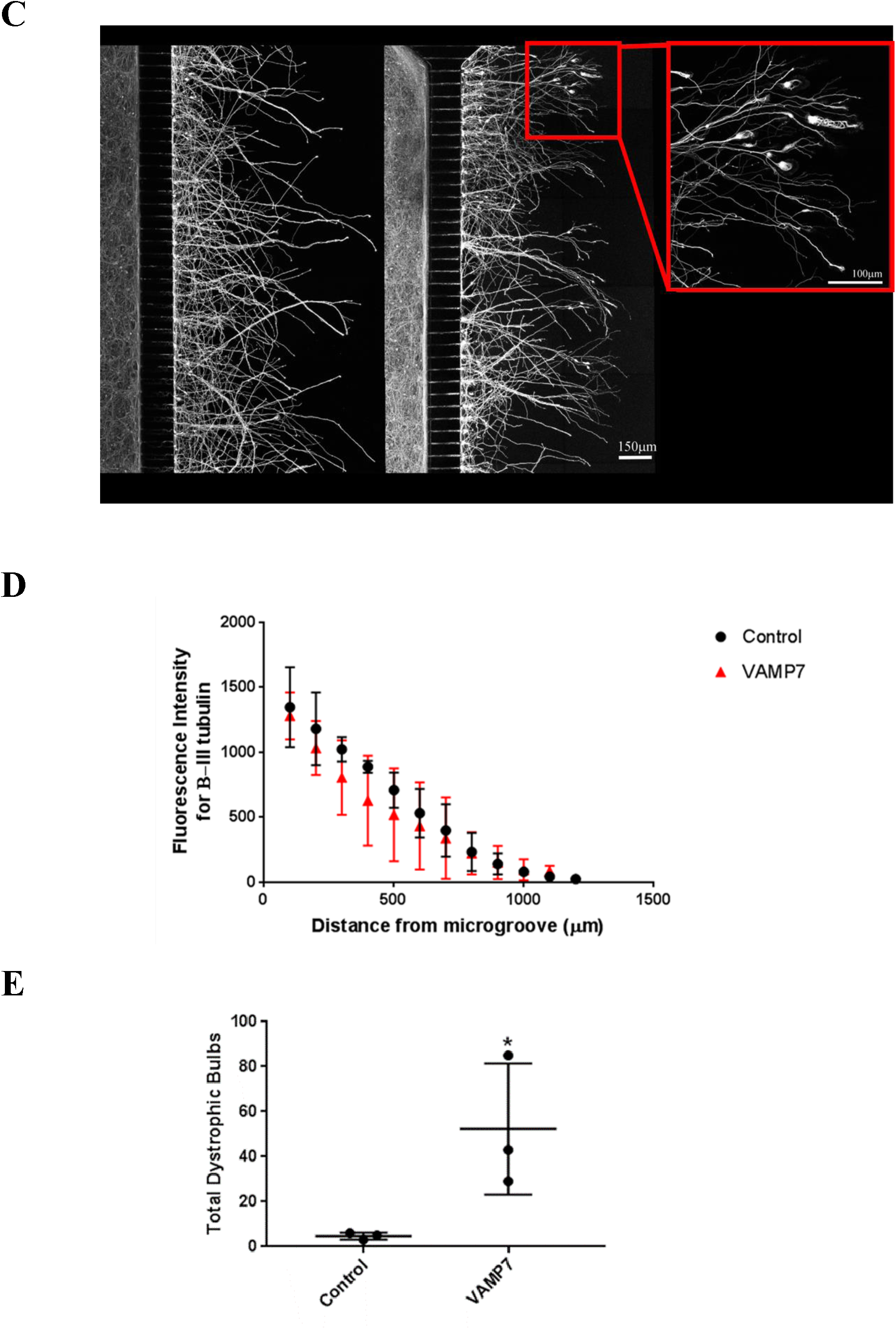
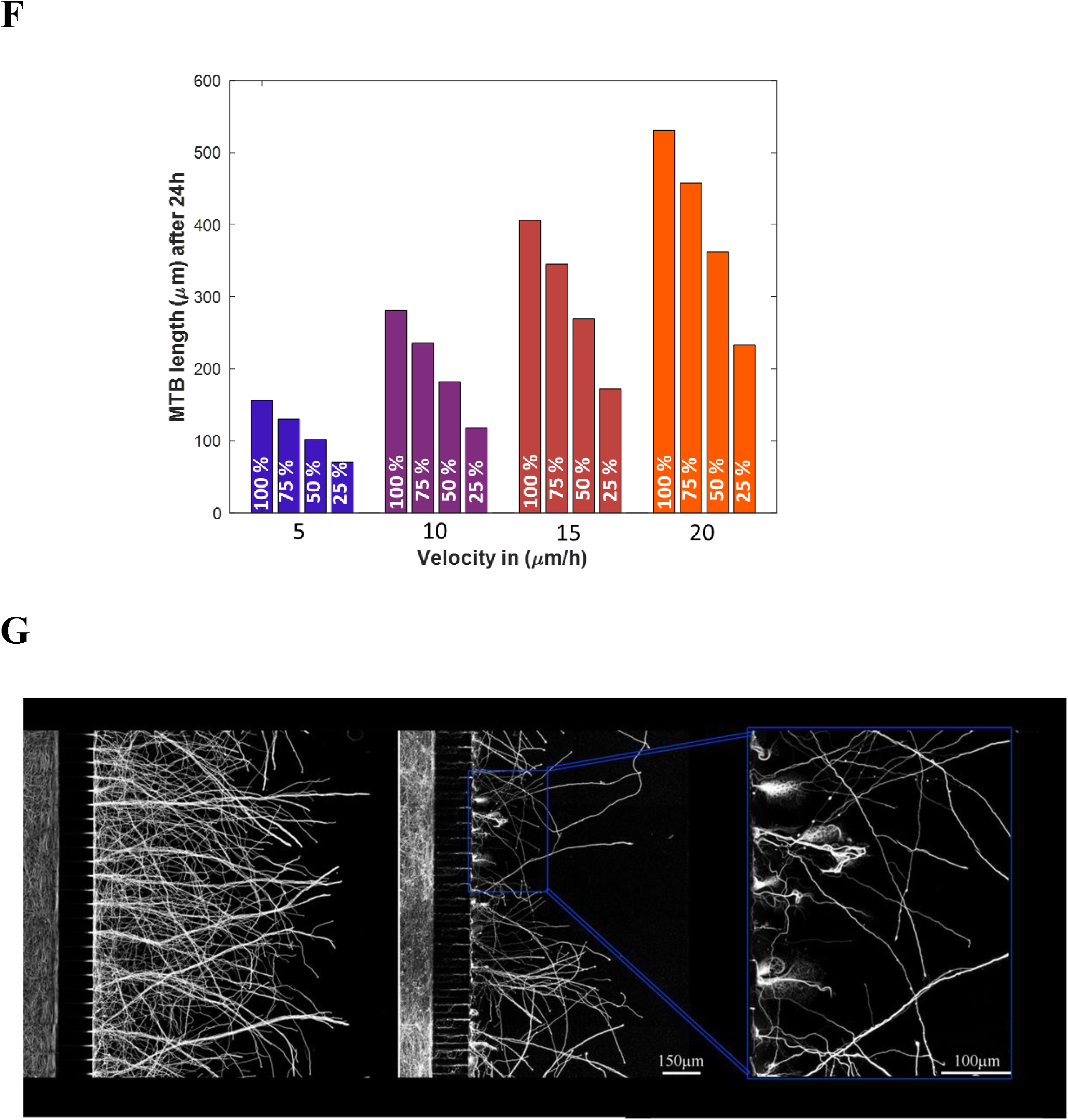
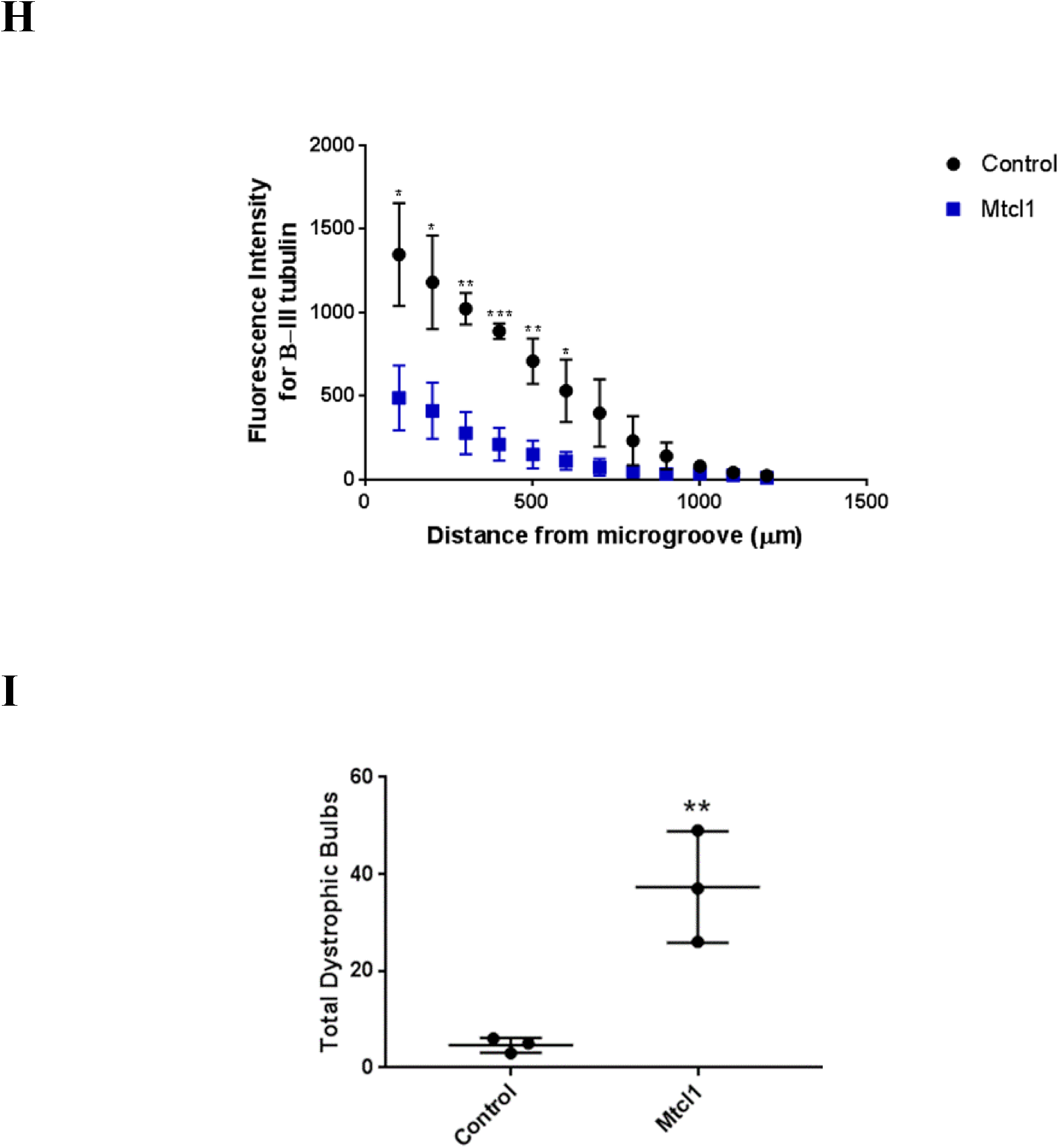
Imbalanced SCP dynamics disturb neurite outgrowth. **(A-B)** We predicted four sets of kinetic parameters that allow coordinated NOG at the velocities 5, 10, 15 and 20 μm/h. To simulate the effect of uncoordinated vesicle SCP dynamics we decreased the count and production rate of v-SNARE V to 25%, 50% and 75% of the original prediction, without modifying any other parameters. Numerical simulations predict that reduced exocytosis causes an increase in the count of vesicles in the growth cone cytoplasm (A) without reducing NOG velocity (B). **(C)** We validated our prediction by analyzing neurite outgrowth after knockdown of the v-SNARE VAMP7. P1 rat cortical neurons transfected with siRNA against VAMP7 or scrambled siRNAs were stained for β-III tubulin to document neurite growth across the microgroove of the chamber. In agreement with our prediction, VAMP7 knock down did not decrease total outgrowth length. It caused a significant number of physiological growth cones to turn into dystrophic bulbs. **(D)** Analysis of quantified neurite lengths confirmed that there was no significant difference in outgrowth length. Shown are average values and standard deviations (three independent experiments). **(E)** Neurites growing from neurons after VAMP7 knock down showed a significant increase in the number of dystrophic bulbs, indicative for impaired NOG. **(F)** To simulate the effect of reduced MT stabilization, we reduced the stabilization rate that turns dynamic into stable MTs to 25%, 50% and 75% of the original prediction, without changing the predicted values for all other kinetic parameters. Numerical simulations propose that decreasing stabilization leads to a significant decrease in NOG outgrowth over the investigated time. **(G)** We validated our predictions by knock down of the microtubule crosslinking protein MTCL1. MTCL1 knock down caused reduced NOG outgrowth and dystrophic bulb formation. **(H)** Outgrowth quantification confirms significant decrease in NOG (3 independent experiments). **(I)** Mtcl1 also significantly increased the number of dystrophic bulbs. *p<0.05, **p<0.01, ***p<0.001based on unpaired t-test for all panels.

To simulate impaired activity of the Level-2 SCP *Stable Microtubule Growth* we selected its Level-3 SCP child *Microtubule Stabilization* and perturbed the kinetic parameter stabilization rate. Our simulations predicted a significant decrease in the total MTB length that affects NOG (Figure 9F). We validated our predictions by knocking down the microtubule crosslinking protein MTCL1 that mediates MT stabilization (Satake et al., 2017). MTCL1 KO significantly reduced NOG velocity and increased the number of dystrophic bulbs (Figure 9G/H/I).

## Discussion

### Whole cell responses depend on subcellular activities at different levels

Neurite outgrowth depends on the coordinated activities of various sub-cellular processes (SCPs). Of the different SCPs involved in our model, the types of Level-1 SCPs are critical. Three major sub-cellular functions that are covered by these SCPs are *Production of Membrane Components, Vesicle Transport & Exocytosis* and *Microtubule Growth*. We curated SCPs that are involved in these Level-1 sub-cellular functions and classified them as Level-2 and Level-3 SCPs that are in parent-child relationships. Based on the hierarchical relationships between these SCPs, we developed a dynamical model that incorporates the three types of Level-1 SCPs and documented that the activity of the SCPs involved need to be highly coordinated to allow neurite outgrowth at a certain velocity. To make the model more manageable, our simulations were done under several assumptions: (1) the size of the TGN & GC-PM (50 μm^2^) is independent of the outgrowth velocity; (2) the rate of back transported membrane from the growth cone to the TGN is 0.5 μm^2^/min; (3) 90 % of the anterograde vesicles in the neurite shaft cytoplasm and the growth cone cytoplasm serve as membrane reservoir (i.e. these vesicles are not actively transported along the MT or fusing with the GC-PM) and (4) 20 μm of the neurite microtubule scaffold consists of dynamic MTs and the rest of stable MTs. All these assumptions are based on previous experimental data and allowed us to identify the key features of the balance between SCP types required for a dynamic whole cell response. We developed an analytical prediction for steady state dynamics at the specified NOG velocities that allowed us to investigate SCP relationships in a systematic manner.

### Complementary dependence in lower level SCPs enables robustness of response, while imbalances in higher level SCPs are not readily compensated

We could show that the activities of Level-3 SCP siblings needed to generate the function of their Level-2 SCP parent inversely depends on each other. This applied to both Level-3 SCPs involved in *Vesicular Transport & Exocytosis* as well as Level-3 SCPs involved in *Microtubule Growth*. This complementary compensatory capability will confer the cell to ability to mount a response in a robust manner

In contrast, the activities of Level-2 SCPs must be highly coordinated to maintain the Level-1 SCP function. The impaired activity of one Level-2 SCP cannot be simply compensated by an increased activity of another Level-2 SCP. Dependencies between Level-2 SCPs are more complex. In case of *Vesicle Transport & Exocytosis*, loss of function of one Level-2 SCP might only be compensated, if multiple other Level-2 SCPs also reduce their function. Impaired Level-2 SCP activities were simulated by reduction of the activity of one of its Level-3 children SCPs without compensatory activity modification of the other.

Our predictions proposed that reduced exocytosis does not influence NOG velocity. We predicted that this results in an excess of anterograde vesicle in the GC that can be one of the reasons for the formation of dystrophic bulbs. In agreement with our prediction, knockdown of v-SNARE VAMP7 did not change NOG velocity but rather, it creates a pathological phenotype with multiple dystrophic bulbs instead of healthy growth cones.

A reduction in the growth of stable MTs was predicted to cause a reduction in NOG length, and it was experimentally confirmed by knock down of MTCL1. As the activities of the SCPs related to *Vesicle Transport & Exocytosis* are unchanged, in this case the reduction in neurite outgrowth will be accompanied by the appearance of dystrophic bulbs. This prediction was also experimentally confirmed.

### Different subcellular processes can give rise to similar whole cell phenotypes

The observations in this study both computational and experimental demonstrate the critical need for balance and coordination between different types of subcellular processes required to mount a whole cell response. It is noteworthy that robustness of whole cell responses arises from the relative redundancy of lower level subcellular processes within each type. However at least for neurite outgrowth, such redundancies do not occur at higher level SCPs where, when imbalance occurs the whole cell function shuts down even though each high level SCP might be operational for a certain time period. It is the uncoordinated functioning of the higher level SCPs that result in the morphological abnormalities like the dystrophic bulbs. Since lack of coordination can arise to due to dysfunction of different higher level SCPs we get similar dystrophic bulbs when we partially ablate a microtubule stabilizing protein or a vesicle fusion protein. This provides a striking example of phenotypic convergence in spite of molecular divergence.

Overall the major conclusions of this study are that 1) elements of both robustness and fragility coexist within subcellular processes needed to produce whole cell responses; 2) since multiple types of subcellular processes are involved in mounting a whole cell responses changes in in different molecular entities or subcellular processes can result in the same aberrant phenotype; 3) quantitative imbalances between different types of subcellular processes are sufficient for alteration in cellular dynamics causing abnormal phenotype.

## Acknowledgements

This research was supported by NIH grant GM54508 and the Systems Biology Center grant GM 071558. We thank Alan Stern and Marc Birtwistle with help in image acquisition using the InCell analyzer and Frank Bradke, Tatiana Stepanova and Ali Erturk for providing experimental data for constraining our model. We thank Nikolaos Tzavaras for helping us in image quantification in MetaMorph. We thank Drs Joseph Goldfarb and Robert D Blitzer for critical reading of the manuscript.

## Author contributions

R.I. and J.H. conceived the project. A.Y. and J.H. developed and implemented the model. J.H. developed the analytical prediction for steady state dynamics. M.S, V.R. and R.T. prepared rat cortical neurons. A.Y. and M.S. analyzed the images. A.Y., R.I. and J.H. wrote the manuscript.

**Declaration of Interests:** The authors declare no competing interests.

